# Overexpression of Ssd1 and calorie restriction extend yeast replicative lifespan by preventing deleterious age-dependent iron uptake

**DOI:** 10.1101/2025.09.02.673772

**Authors:** J. Ignacio Gutierrez, Claudia Edgar, Jessica K. Tyler

**Affiliations:** Department of Pathology and Laboratory Medicine, Weill Cornell Medicine, 1300 York Avenue, New York, NY 10065, USA; BCMB Graduate Program, Weill Cornell Graduate School of Biomedical Sciences, New York, NY 10065, USA

## Abstract

Overexpression of the mRNA binding protein Ssd1 extends the yeast replicative lifespan. Using microfluidics to trap and image single cells throughout their lifespans, we find that lifespan extension by Ssd1 overexpression is accompanied by formation of cytoplasmic Ssd1 foci. The age-dependent Ssd1 foci are condensates that appear dynamically in a cell cycle-dependent manner and their failure to resolve during mitosis coincided with the end of lifespan. Ssd1 overexpression was epistatic with calorie restriction (CR) for lifespan extension and yeast overexpressing Ssd1 or undergoing CR were resistant to iron supplementation-induced lifespan shortening while their lifespans were reduced by iron chelation. The nuclear translocation of the Aft1 transcriptional regulator of the iron regulon occurred during aging in a manner that predicted remaining lifespan, but was prevented by CR. Accordingly, age-dependent induction of the Fit2 and Arn1 high-affinity iron transporters within the iron regulon was reduced by CR and Ssd1 overexpression. Consistent with age-dependent activation of the iron regulon, intracellular iron accumulated during aging but was prevented by CR and Ssd1 overexpression. Moreover, lifespan extension by Ssd1 overexpression or CR was epistatic to inactivation of the iron regulon. These studies reveal that CR and Ssd1 overexpression extend the yeast replicative lifespan by blocking deleterious age-dependent iron uptake, identifying novel therapeutic targets for lifespan extension and providing insight into how CR may extend the lifespan and healthspan in humans.

## Background

Aging is a gradual and irreversible biological process that results in the progressive decline in physical function of cells and tissues. The fundamental processes that go awry during aging are conserved across eukaryotes, exemplified by the fact that anti-aging regimens such as calorie restriction (CR) and mTOR/Tor1 pathway inhibition using rapamycin function to extend lifespan in species ranging from budding yeast to mammals^1,2^. There is a great deal of interest in the identification of common features, or hallmarks, of aging^3^ and in determining which of these hallmarks are reversed by anti-aging regimens.

Loss of cellular homeostasis is a hallmark of aging, characterized by the gradual decline in the ability to regulate intracellular conditions leading to accumulation of cellular damage, impaired stress responses, and functional deterioration across tissues^3,4^. Age-associated dysregulation of transcription and translation may contribute to this loss of homeostasis during aging as it can cause imbalanced protein production, impaired stress responses, and metabolic disruption leading to accumulation of damaged proteins^5,6^. In this context, Ssd1, a yeast mRNA-binding protein involved in modulating the stability, localization, or translation of specific mRNAs^7^, may play a pivotal role in preserving cellular homeostasis during aging given that its deletion shortens the yeast replicative lifespan^8^ while its overexpression extends lifespan^6^. Yeast is the only type of eukaryotic cell where the length of the replicative lifespan (RLS), the number of times that a cell divides before death, can be measured accurately^9^. Previously, we have shown that overexpression of Ssd1 extends yeast RLS by suppressing global translation efficiency^6^ possibly involving its stress-induced role in mRNA sequestration into cytoplasmic mRNA storage granules termed P-bodies and stress granules where the mRNAs can be stored or degraded^10^. Ssd1 has been shown to bind a large number of mRNAs, with some reports identifying up to 300 targets^11,12^. However, whether sequestration of a subset of these mRNAs by overexpressed Ssd1 is sufficient for lifespan extension remains unclear. Additionally, the identity of the mRNAs whose translation is suppressed by overexpressed Ssd1 to extend lifespan is unknown.

By contrast to yeast lifespan extension by Ssd1 overexpression, calorie restriction (CR) has been shown to extend lifespan in many organisms including primates^13,14^. However, CR reprograms metabolism affecting a large portion of cellular processes^1^, making it difficult to determine the precise molecular mechanism of CR-mediated lifespan extension. One putative mechanism for improved cellular homeostasis by CR leading to lifespan extension implicates CR’s ability to reduce reactive oxygen species (ROS) production, thereby decreasing oxidative stress^15^. Studies in yeast have suggested that CR reduces ROS levels by mechanisms including activating superoxide dismutases^16^ as well as by improving mitochondrial function^17,18^. Another mode for reduction of oxidative stress by CR, at least in the mouse heart, is decreased cellular iron uptake to improve iron homeostasis^19^. In this situation, mechanistically CR resulted in the downregulation of transcription of genes encoding iron transporters that mediate iron import and upregulation of ferritin to sequester iron^19^. Whether CR downregulates iron uptake to lead to lifespan extension is an open question.

Iron is an essential cofactor for numerous enzymatic processes, being required for the proper function of enzymes involved in oxygen transport, electron transfer, DNA replication and repair, and various metabolic pathways^20^. However, in its free form (free labile iron, Fe^+2^), iron is highly reactive and induces the production of ROS, thereby generating oxidative stress^21^. For this reason, iron uptake is a tightly regulated process such that when the levels of cellular iron are sufficient, iron transport into the yeast cells is mediated by only low-affinity iron transporters^22^. When iron levels in the cell are deficient, the transcription factors Aft1 and Aft2 translocate from the cytoplasm to the nucleus in yeast^23^ to induce the iron regulon: a set of genes encoding machinery that coordinates iron uptake (including high-affinity iron transporters and transporters of siderophores that scavenge iron from the extracellular environment), vacuolar mobilization, and metabolic adaptation to maintain cellular iron homeostasis^24,25^. Recent reports indicate that the activation of the iron regulon occurs during yeast replicative aging as a result of reduced levels of mitochondrial iron-sulfur clusters^26,27^. We hypothesized that age-dependent activation of the iron regulon would lead to accumulation of free labile iron in cells during aging that would be detrimental to lifespan.

Here we sought to identify the molecular mechanisms whereby Ssd1 overexpression and CR extend the yeast RLS. Using fluorescent imaging of single cells immobilized in microfluidic devices throughout their lifespans, we find that lifespan extension by overexpression of Ssd1 is accompanied by formation of cytoplasmic Ssd1 foci. These foci appear dynamically after DNA replication and resolve by cytokinesis, while their failure to resolve by cytokinesis coincides with the end of lifespan. Furthermore, Ssd1 overexpression is epistatic with CR for lifespan extension. Our results indicate that both CR and Ssd1 overexpression prevent the activation of the iron regulon during aging, which in turn prevents the increased uptake of iron that usually occurs during aging. Furthermore, we show that derailing age-dependent activation of the iron regulon is the molecular mechanism whereby CR and Ssd1 extend replicative lifespan.

## Methods

### Yeast strains

All strains were derived from BY4741^28^. *SSD1* was replaced by a NatMX cassette (FLY2184, gift from Cornelia Kurischko)^10^. To generate *SSD1*-eGFP fusions, we integrated PCR products amplified from plasmid FLE1019^10^ (pAG415-P_GPD1_-*SSD1*) and pFA6a-GFP(S65T)-KanMX6^29^ via homologous recombination. These constructs included the *GPD1* or *SSD1* promoter, respectively and were integrated into the genome at the *SSD1* locus by homologous recombination.

Gene deletions and the addition of the remaining fluorescent tags (except for P_GPD1_-GFP-*AFT1*) were generated by homologous recombination with PCR products from plasmids developed by Pringle et al.^29^. The P_GPD1_-GFP-*AFT1* strain was constructed by homologous recombination of a PCR product from a pRS plasmid containing P_GPD1_-eGFP^30^. All strains used in this study are cataloged in Supplemental Table 1.

### Replicative lifespan analyses

Replicative lifespan (RLS) analyses were performed as follows. Unless otherwise indicated, cells were grown overnight in Synthetic Complete Media (SCD) composed of yeast nitrogen base without amino acids and ammonium sulfate (Difco 233520), complete drop-out mix without YNB (US Biological D9515), ammonium sulfate (Fisher Chemical A702-3), and glucose (Sigma-Aldrich G8270). Cultures were grown to an optical density (OD)_600_ between 0.2 and 0.6, then diluted to an OD_600_ of 0.1, vortexed for 30 seconds, and loaded into microfluidic (described below in more detail) and as we recently described^31^. For experiments involving artemisinin (Art) supplementation, cells were cultured in yeast peptone dextrose (YPD) media containing the indicated concentration of dextrose (Grainger 31GE61; 31GC58), with artemisinin (Thermo Scientific J65406.03) added directly to fresh YPD prior to sterilization. RLS was determined using microfluidic chips from iBiochips (https://ibiochips.com/product/automated-dissection-screening-chip/) according to the manufacturer’s instructions. Where indicated, iron(III) chloride (Sigma 157740-100G) and bathophenanthrolinedisulfonic acid disodium salt hydrate (BPS; Sigma 146617-1G) were added directly to SCD media before sterilization using a Corning filter bottle system with a 0.22 µm pore 54.5 cm² PES membrane (Corning 431098), at the specified final concentrations. All experimental conditions and compound concentrations are detailed in the relevant figure legends.

Each strain was analyzed in the microfluidic chamber under non-fluorescent conditions in three independent experiments, each yielding consistent results, with 30 to 50 cells counted per condition or strain in each experiment. Cells that were lost from the microfluidic traps during the course of the experiment were recorded as censored. For strains expressing fluorescent markers, experiments were conducted under the appropriate fluorescent illumination, also in three independent replicates with consistent results, with 20 to 40 cells counted per experiment. For these fluorescent strains, the total number of cells from all three experiments is presented in the histograms, and only cells that completed their full replicative lifespan were included in the analysis, with cells lost from the traps excluded. Replicative lifespan survival curves were generated using the OASIS 2 platform (https://sbi.postech.ac.kr/oasis2/) as described by Han et al.^32^, and p-values for RLS curves were calculated using the log-rank test.

### Flow cytometry

The settings used on an LSR II flow cytometer were for detection of FITC (excitation 494, emission 520) and TxRed (excitation at 595 nm, emission at 613 nm). For each sample, data from at least 10,000 events were collected, and fluorescence intensity was quantified using pulse area (integral) measurements.

### Image collection

All imaging in this study, including replicative lifespan (RLS) experiments, was performed using an EVOS II microscope (Thermo Fisher). For colocalization experiments, Ssd1-GFP was imaged using the EVOS Light Cube GFP 2.0 (Invitrogen Thermo Fisher, AMEP4951) with light intensity set to 0.053, exposure time to 0.047 seconds, and gain to 1. Hsp104-mCherry, Edc3-mCherry, and Pab1-mCherry were imaged using the EVOS Light Cube TxRed 2.0 (Invitrogen Thermo Fisher, AMEP4955) with light intensity at 0.6, exposure time at 0.06 seconds, and gain at 5. For RLS experiments without fluorescence microscopy, imaging was performed at 40× magnification on V-shaped chips (iBiochips)^31^, with images acquired every 20 minutes for 96 hours. Brightfield illumination was adjusted individually for each experiment, and z-stacks were collected over a physical length of 19 μm in 5 steps of 3.8 μm each. When RLS was measured in conjunction with fluorescence microscopy, imaging was conducted at 60× magnification with immersion oil on Y-shaped AD chips (iBiochips)^31^, with images taken every 20 minutes for 96 hours. Ssd1-GFP was imaged as above, but with an exposure time of 0.045 seconds. Hsp104-mCherry, Edc3-mCherry, and Pab1-mCherry were imaged with light intensity at 0.3, exposure time at 0.06 seconds, and gain at 5. Fit2-mCherry and Arn1-mCherry were imaged with light intensity at 0.105, exposure time at 0.05 seconds, and gain at 4. Aft1-GFP was imaged with light intensity at 0.053, exposure time at 0.04 seconds, and gain at 1. For measurements involving the GFP channel in PGFL experiments, the same light cube was used with light intensity at 0.053, exposure time at 0.07 seconds, and gain at 1. In general, higher exposure times and lower light intensities were preferred to minimize phototoxicity, and all media used for fluorescence imaging was supplemented with 20 mM ascorbic acid (LabChem LC115309) to reduce photobleaching. Fluorescence imaging experiments used z-stacks spanning 8.5 μm in 5 steps of 1.7 μm. All imaging parameters were optimized to achieve the desired temporal resolution while minimizing phototoxicity.

### Image analysis and data representations

The number of cell divisions in each RLS was determined manually by analyzing images captured at 40× magnification every 20 minutes, using the image sequence function in ImageJ. Fluorescence microscopy images were also analyzed manually in ImageJ, with appropriate background corrections applied. For Ssd1 foci quantification, a minimum threshold of 200-pixel intensity above background was used to score foci; structures below this threshold were disregarded. Multiple foci per cell cycle were recorded if two or more foci were observed simultaneously in the same cell, or if a focus disappeared and a new focus appeared in subsequent images within the same cell cycle. For single-cell trajectory histograms, individual values for fluorescence intensity or foci number, depending on the experiment, were compiled from three independent experimental repeats and visualized in Excel by replacing values with the corresponding color codes in the cells. Dot plots were generated using Prism, while box-and-whisker plots were created in Excel. Statistical significance for dot plots and box-and-whisker plots was assessed using the two-sided Student’s t-test. All analyses were performed in at least three independent experiments with reproducible results, and the combined data are presented in the figures.

### Iron measurements

Intracellular free iron (Fe^+2^) levels were measured using Phen Green™ FL, Diacetate, cell permeant (Thermo Fisher P6763), following a yeast-adapted protocol as described by Gomez-Gallardo et al^33^. Briefly, cells were aged in the microfluidic chamber as previously described, and at approximately 30 hours, the chip was loaded with 5 µg/mL Phen Green FL in DMSO. GFP fluorescence was measured using the EVOS Light Cube GFP 2.0 (Invitrogen Thermo Fisher, AMEP4951). The chip was then loaded with 1 mM of the iron chelator 1,10-phenanthroline (Sigma 131377-5G), and fluorescence was measured again. The ratio of fluorescence before and after chelator addition is proportional to the concentration of intracellular free labile iron in the cell.

## Results

### Lifespan extension by overexpression of Ssd1 is accompanied by age-dependent appearance of Ssd1 condensates

The Ssd1 mRNA-binding protein has a prion-like domain^34^ which has been shown to be involved in the localization of overexpressed Ssd1 to cytoplasmic condensates including P-bodies and stress granules in response to glucose starvation^34^. Because the conditions in which Ssd1 overexpression extends the yeast lifespan are glucose replete^6^, we asked whether Ssd1 even forms foci / condensates during replicative aging. Therefore, we generated a yeast strain overexpressing eGFP-tagged Ssd1 by integrating *SSD1*-eGFP under the control of a strong *GPD1* promoter (P_GDP1_-SSD1-GFP). This promoter has been used extensively to overexpress *SSD1* in previous studies^10,34^. We also made a strain expressing eGFP-tagged Ssd1 from the endogenous *SSD1* promoter (SSD1-GFP). We confirmed by flow cytometry that overexpression of Ssd1 occurred in seemingly all P_GDP1_-SSD1-GFP cells (**Figure 1A**). We then measured the RLS of these strains on microfluidic devices^31^ and observed a robust lifespan extension in the strain overexpressing *SSD1* (P_GPD1_-SSD1-GFP) (**Figure 1B**). By contrast, a strain deleted for *SSD1* had a shortened RLS (**Figure 1B**), as published previously^8^.

**Figure 1.**
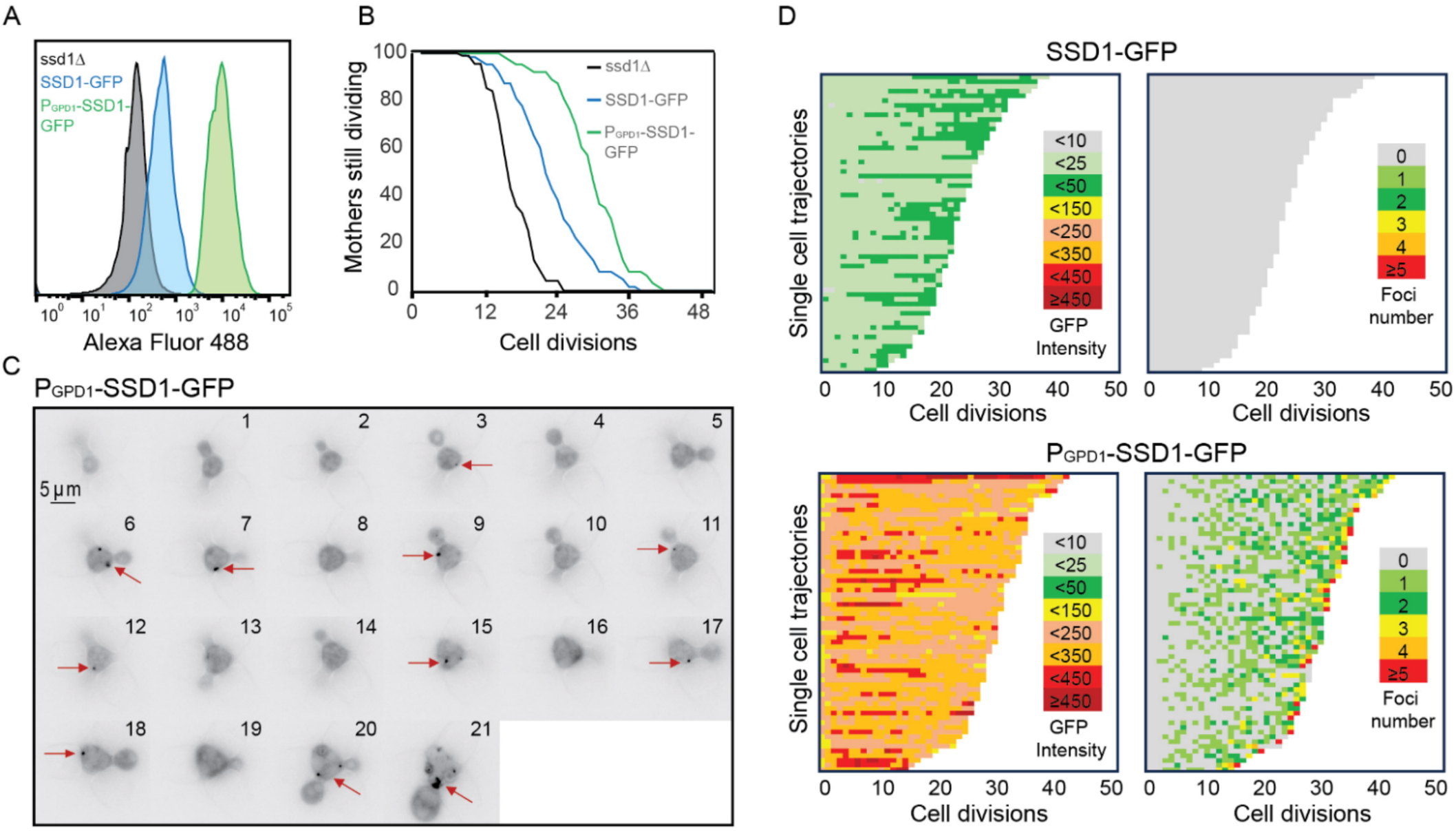
Overexpression of Ssd1 extends lifespan and forms transient cytoplasmic foci during replicative aging. **A.** Levels of GFP tagged Ssd1 in the indicated strains, for 10000 cells per strain. **B** RLS of strains shown in panel A for 90 *ssd1Δ* cells, 63 SSD1-GFP cells and 63 P_gpd1_-SSD1-GFP cells, *p-* values are as follows: for control strain versus ssd1Δ strain is 0, for control versus P_gpd1_-SSD1-GFP is 8 x 10^-8^ **C.** Transient formation of Ssd1-GFP foci in P_gpd1_-SSD1-GFP strain during aging at the indicated cell division number of the RLS for one cell. Arrows indicate Ssd1-GFP foci. **D.** Single cell trajectories showing total cell Ssd1-GFP intensity and number of Ssd1-GFP foci formation for the indicated strains throughout the RLS. Each rectangle represents one cell cycle.

We next asked whether overexpression of Ssd1 leads to formation of Ssd1 foci during replicative aging. Using microfluidics combined with fluorescence microscopy^31^, we tracked single cells from the SSD1-GFP (control) or P_GDP1_-SSD1-GFP strains throughout their entire lifespan, acquiring images every 20 minutes over approximately 4 days. We observed that in P_GPD1_-SSD1-GFP cells, but not in the control strain, transient Ssd1 foci appeared during aging (**Figure 1C and 1D**, **Figure 1-figure supplement 1, Supplemental Movie 1**). The abundance of the Ssd1 protein increased over the first 4–6 cell divisions of the RLS before reaching a plateau, both in control and P_GPD1_-SSD1-GFP cells (**Figure 1D**), where the overall fluorescence intensity in P_GPD1_-SSD1-GFP cells was approximately tenfold higher than in the control strain (**Figure 1D, Sup. Figure 1-figure supplement 2**). No Ssd1-GFP foci were observed in P_GPD1_-SSD1-GFP cells during the early cell divisions in the lifespan (**Figure 1D**), suggesting that either a threshold concentration of Ssd1 may be required for foci formation and / or that alterations in cell homeostasis during aging promote Ssd1 foci formation in cells overexpressing Ssd1. During aging, Ssd1 foci appeared in all cells overexpressing Ssd1, and the number and frequency of Ssd1 foci appearance varied between cells, with the highest number of foci per cell apparent at the end of the lifespan (**Figure 1D**). These data show that the lifespan extension that results from overexpression of Ssd1 is accompanied by age-dependent formation of Ssd1 foci, suggesting that the ability to form Ssd1 foci correlates with increased longevity while the appearance of five or more Ssd1 foci may be either a cause or consequence of end of lifespan.

Similar to the starvation-induced Ssd1-GFP foci in Ssd1 overexpressing cells^34^, age-induced Ssd1-GFP foci in Ssd1 overexpressing cells are also cytoplasmic (**Figure 1-figure supplement 3A**). Moreover, Ssd1-GFP foci can be dissolved by 1,6, hexanediol suggesting that they are condensates (**Figure 1-figure supplement 3B**). Furthermore, age induced Ssd1-GFP foci underwent fusion and fission (**Supplemental Movie 2**), consistent with them also being condensates.

### Ssd1 condensates form in cells overexpressing Ssd1 during aging in a cell cycle regulated manner and are distinct from age-induced P-bodies and Hsp104 foci

To investigate further the transient nature of appearance of the Ssd1 condensates during aging (**Figure 1C and 1D**), we used bud size as a proxy for cell cycle stage (**Figure 2A**). We found that Ssd1 condensates typically appeared in cells overexpressing Ssd1 during aging in the G_2_/M cell cycle stages, when the bud reaches its maximum size, and very rarely during G_1_/S, when the bud was absent or small (**Figure 2A**). We also observed that Ssd1 condensates virtually always lasted less than one cell cycle, dissolving before or during cytokinesis of the same cell cycle in which they formed (**Figure 2A**). However, the Ssd1 condensates were not dissolved before mitosis in some cells in their last division of their RLS (**Figure 2A**). As such, it appears that the ability to both form and resolve Ssd1 foci during aging correlates with lifespan extension, while failure to resolve Ssd1 foci correlates with the end of lifespan in some cells.

**Figure 2.**
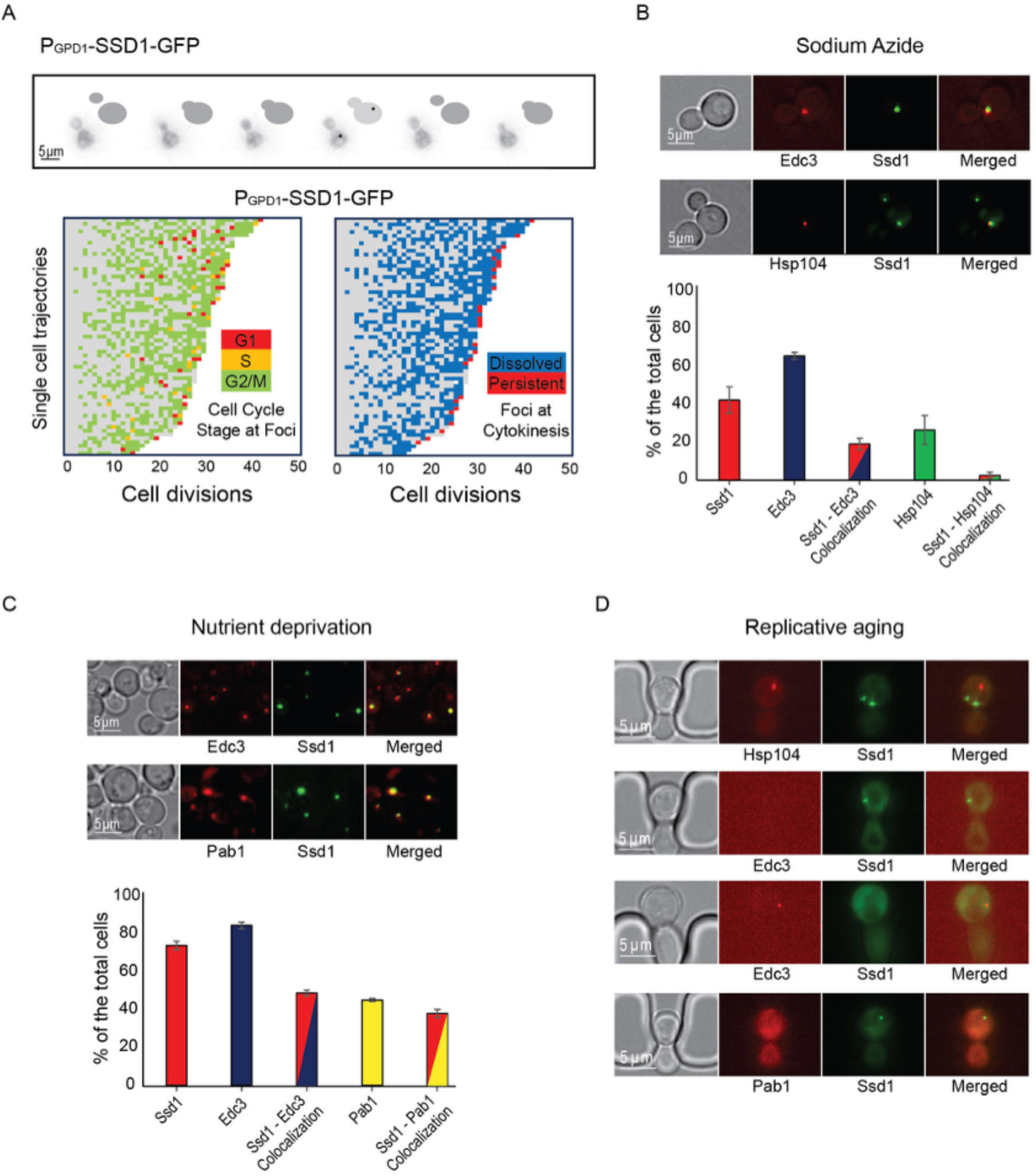
Ssd1 foci formed during aging are cell cycle regulated and distinct from Hsp104 foci, P-bodies and stress granules. **A.** Example showing Ssd1-GFP foci in G_2_ phase during aging. The images are taken of the same cell every 20 minutes and the cartoons indicate the bud size. Below are shown single cell trajectories of cell cycle stage at which Ssd1 foci are present during aging (left) and whether the foci dissolved at the following cytokinesis (shown in blue) or were persistent through cytokinesis (shown in red) (right). **B.** Colocalization of Ssd1 foci with P-bodies and Hsp104 foci in cells treated with 10 mM Sodium azide. Quantitation is shown below for over 100 cells per condition. **C.** Colocalization of Ssd1 foci with P-bodies (marked by Edc3) and stress granules (marked by Pab1) under nutrient deprivation and quantitation. **D.** Failure of Ssd1 foci to colocalize with age-induced Hsp104 foci or P-bodies.

We were curious to investigate the relationship of the Ssd1 condensates that occur in cells overexpressing Ssd1 during aging to other cytoplasmic condensates that Ssd1 has been reported to colocalize with, when overexpressed. Ssd1 foci formation has primarily been studied in the context of the stress response^34^, where overexpressed Ssd1 colocalizes with stress granules and P-bodies upon glucose depletion, hypertonic stress, and heat shock, for translational repression by sequestering untranslated mRNAs^10^, or helping mitigate proteotoxic stress by directing misfolded proteins to IPODs (insoluble protein deposit sites)^34,35^. While the strain overexpressing Ssd1 (P_GPD1_-SSD1) only very rarely exhibited Ssd1 foci during exponential growth (**Figure 2-figure supplement 1**), we observed a significant increase in Ssd1 foci formation under stress: approximately 40% and 75% of P_GPD1_-SSD1 cells displayed Ssd1 foci following treatment with 10 mM sodium azide or growth to saturation (nutrient deprivation), respectively (**Figure 2B, 2C**). Sodium azide, which inhibits the electron transport chain, induced P-body formation in ∼65% of cells, as marked by mCherry-tagged Edc3. Approximately 20% of cells showed Ssd1 localized to P-bodies in response to sodium azide treatment (**Figure 2B**). Similarly, tagging Hsp104 with mCherry revealed that upon sodium azide treatment ∼25% of cells showed Hsp104 foci and 5% of cells had Hsp104 colocalizing with Ssd1 foci (**Figure 2B**). Under nutrient deprivation, ∼85% of cells formed P-bodies and ∼50% of cells showed colocalization with Ssd1 foci (**Figure 2C**). In the same condition, ∼45% of cells exhibited stress granules, as indicated by mCherry-tagged Pab1 and ∼40% of cells also displayed colocalization with Ssd1 foci (**Figure 2C**). These results indicate that in cells overexpressing Ssd1, Ssd1 frequently but not always, colocalized with stress-induced foci under acute stress conditions.

To determine whether Ssd1 foci that form during aging also colocalize with P-bodies, stress granules or Hsp104 foci, we examined these condensates in aging P_GPD1_-SSD1 cells. Hsp104 foci have been shown to form around the fifth cell division of the RLS and are asymmetrically retained in mother cells throughout the lifespan^36^. Consistent with this, we observed Hsp104 foci during aging, but we never observed them colocalized with Ssd1 condensates (**Figure 2D**) even though Ssd1-GFP condensates form in every cell during aging (**Figure 1D**). Previous reports indicate that P-bodies appear in late lifespan^6^; we similarly observed Edc3-mCherry foci (marking P-bodies) in late lifespan, but again, we did not observe them ever colocalizing with Ssd1 foci (**Figure 2D**). We did not observe the formation of stress granules as determined by Pab1-mCherry staining during the RLS (**Figure 2D**). These data indicate that the Ssd1 condensates that form during aging in cells overexpressing Ssd1 are distinct from P-bodies, stress granules and Hsp104 foci.

### Overexpression of Ssd1 is epistatic to calorie restriction for lifespan extension and both reduce activation of the iron regulon during aging

To gain mechanistic insight into how overexpression of Ssd1 and Ssd1 condensate formation may extend lifespan, we tested whether overexpression of Ssd1 (P_GPD1_-SSD1) was epistatic to CR for lifespan extension. We found that the RLS extension resulting from Ssd1 overexpression was equivalent to that achieved by CR (0.05% glucose) and moreover, was equivalent to the RLS of cells that overexpressed Ssd1 during CR (**Figure 3A**). These results are consistent with Ssd1 overexpression and CR extending lifespan via a shared pathway.

**Figure 3.**
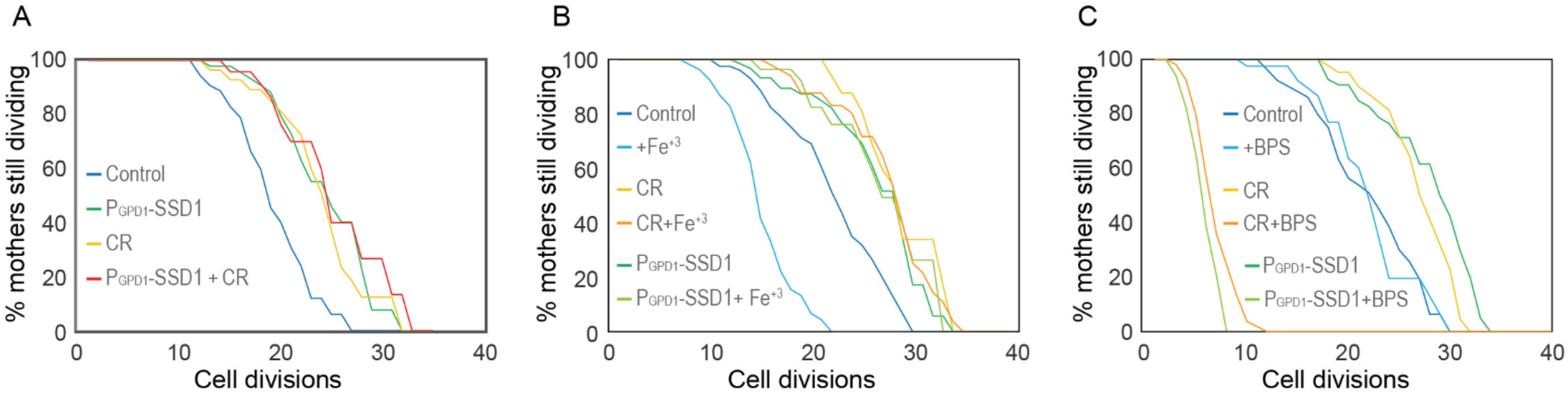
CR and overexpression of Ssd1 are epistatic for lifespan extension, both influencing iron metabolism. **A.** Calorie restriction and overexpression of Ssd1 are epistatic for extension of RLS, for 54 control cells, 48 P_gpd_-SSD1 cells, 48 control cells undergoing CR, and 44 P_gpd_-SSD1 cells + CR. p-values determined by student’s T-test for control versus P_gpd_-SSD1, CR and P_GPD_-SSD1 + CR are 1 x 10^6^, 3 x 10^-4^ and 5 x 10“* respectively. There is no significant difference between P_gpd_-SSD1, CR and P_gpd_-SSD1 + CR. **B.** CR and overexpression of Ssd1 protect cells from lifespan reduction due to iron supplementation with 10 µM Fe^+3^, for 55 control cells, 53 control cells + Fe^+3^, 47 P_gpd_-SSD1 cells, 41 P_gpd_-SSD1 + Fe^+3^ cells, 37 CR cells, and 50 CR + Fe^+3^ cells. p-value for control versus Fe^+3^ is 6 × 10^-7^. There is no significant difference between P_gpd_-SSD1 with and without Fe^+3^ or for CR with and without Fe’^3^. **C.** CR and Ssd1 overexpression protect cells from the lifespan reduction caused by iron depletion with 200 µM BPS, for 52 control cells, 40 control cells + BPS, 60 P_gpd_-SSD1 cells, 34 P_gpd_-SSD1 cells + BPS, 44 control cells under CR, and 53 cells under CR + BPS. p-values for the control + BPS, versus P_gpd_-SSD1 + BPS and CR +BPS are 0, while there is no significant difference between the control with and without BPS treated and untreated control are not significantly different, control compared to P_gpd_-SSD1 and CR are 0.

Other studies in the lab on the effect of CR on iron metabolism had led us to be interested in the observation that the iron regulon is activated during aging^26,27^, presumably resulting in increased iron uptake in aged cells, which is likely to be deleterious for lifespan given that constitutive activation of the iron regulon shortens RLS^25^. As such, we were curious whether overexpression of Ssd1 or CR may lead to a muted iron response during aging. If this were the case, we would expect the lifespan of cells overexpressing Ssd1 or undergoing CR to be less sensitive to iron supplementation. Indeed, the control strain had a significantly reduced RLS upon supplementation with 10µM iron(III) chloride (FeCl₃), while the extended RLS of P_GPD1_-SSD1 and CR-treated cells was unaffected by 10 µM FeCl₃ (**Figure 3B**). To further probe iron dependency, we supplemented the media with 200 µM of the iron chelator bathophenanthroline disulfonic acid (BPS)^37^. The RLS of control cells was unaffected by BPS, whereas P_GPD1_-SSD1 and CR-treated cells exhibited a dramatically shortened lifespan (**Figure 3C**). These data demonstrate that yeast overexpressing Ssd1 or undergoing CR extend RLS by a shared pathway that is resistant to iron supplementation but sensitive to iron chelation.

### Overexpression of Ssd1 and calorie restriction derail the iron regulon during aging

Given that the lifespan extension caused by overexpressing Ssd1 and CR was not prevented by iron supplementation, which shortened the lifespan of control cells (**Figure 3B**), we sought to determine whether CR or Ssd1 overexpression disrupted age-dependent activation of the iron regulon which could theoretically prevent deleterious iron uptake. Under low-iron conditions Aft1 translocates to the nucleus to activate transcription of the iron regulon^23^ (**Figure 4A**, **Figure 4 - figure supplement 1A**). We asked whether Aft1 localizes to the nucleus during aging and whether this is prevented in yeast overexpressing Ssd1 or during CR. Due to its low endogenous expression, we overexpressed a GFP-tagged version of *AFT1* from the GPD1 promoter, as previously reported^38^, to more effectively detect its cellular location. In control cells, we observed nuclear localization of GFP-Aft1 during mid-to-late lifespan (**Figure 4B**). Furthermore, there is a clear trend apparent where cells in which Aft1 translocated to the nucleus later in their lifespan tended to live longer (**Figure 4C**). Also, cells lived on average 5 more divisions after Aft1 translocated into the nucleus (**Figure 4-figure supplement 1B**) suggesting that Aft1 nuclear translocation may predict the end of lifespan. By contrast, CR-treated cells showed little or no nuclear localization of Aft1, except in some cells where it occurred very late in lifespan (**Figure 4B**). Notably, this is not due to reduced levels of Aft1 during CR (**Figure 4-figure supplement 1C**). These data indicate that the iron regulon is activated during aging and is predictive of the end of lifespan, while cells undergoing CR do not activate the iron regulon during aging.

We were unable to examine Aft1 localization during aging in the strain overexpressing Ssd1 because the high-level expression of GFP-Aft1 in this strain caused synthetic sickness. Instead, we sought to examine whether Ssd1 overexpression or CR prevent the induction of gene products within the iron regulon during aging. We analyzed expression of *FIT2* and *ARN1*, tagged with mCherry, two previously used reporters of iron regulon activity^39,40^. Since loss-of-function mutations in *FIT2* and *ARN1* have been shown to extend the RLS^41–43^, we first confirmed that tagging these genes did not impair function: both control and P_GPD1_-SSD1 strains carrying either mCherry tag exhibited an RLS equivalent to the untagged proteins under glucose rich and CR conditions (**Figure 4-figure supplement 2, Figure 3A**). In control strains, expression of Fit2 and Arn1 varied across the population, but generally increased with age (**Figure 4D, 4E**, **Figure 4- figure supplement 3**). In the strains overexpressing Ssd1 fewer cells induced Fit2 and Arn1 during aging compared to control strains, and their overall expression levels were lower upon Ssd1 overexpression (**Figure 4D, 4E**). In CR-treated strains, Fit2 and Arn1 levels remained very low throughout the lifespan (**Figure 4D, 4E**). Quantification of maximum expression per cell confirmed significantly higher levels of Fit2 and Arn1 in the control strain compared to P_GPD1_-SSD1 and CR-treated cells (**Figure 4D, 4E**). These results demonstrate that overexpression of Ssd1 and CR greatly reduce age-related activation of the iron regulon.

It has been reported that preventing activation of the iron regulon benefits longevity by reducing Cth2 expression (*CTH2* is within the iron regulon), an mRNA-binding protein that targets for degradation the mRNA of non-essential iron binding proteins^27^. We deleted *CTH2* and in accordance with the literature, it extended lifespan compared to control strains, however to a much lower extent than overexpressing Ssd1 or CR (**Figure 4-figure supplement 4**). This result indicates that preventing activation of the iron regulon benefits longevity in a manner that is not only due to preventing Cth2 expression.

### Experimental activation of the iron regulon prevents CR or Ssd1 overexpression from extending lifespan

To directly test whether activation of the iron regulon is a negative regulator of longevity, we examined the RLS in strains with a pre-activated iron regulon. In our experiments with the iron chelator BPS, we observed that while P_GPD1_-SSD1 and CR-treated strains exhibited shorter lifespans in media supplemented with BPS, those strains were still capable of overnight growth in the presence of BPS. We took advantage of this observation by growing strains carrying the *FIT2*- mCherry or *ARN1*-mCherry reporter in media supplemented with 200 μM BPS overnight. Flow cytometry analysis of these exponentially growing cultures confirmed strong induction of the iron regulon in the control strain, the strain overexpressing Ssd1 and during CR (**Figure 5A**). We then measured RLS in the iron regulon “induced” populations by transferring them to fresh control or CR media without BPS for the RLS measurements. Only cells showing *FIT2*-mCherry expression (i.e., those with an induced iron regulon) were tracked and compared to their counterparts grown without prior BPS exposure. We found that activation of the iron regulon reduced the RLS of the strain overexpressing Ssd1 and undergoing CR to a level equivalent to the RLS of the control strain (**Figure 5B**). As such, prior activation of the iron regulon prevents CR and Ssd1 overexpression from extending lifespan. These results are consistent with a model where suppression of activation of the iron regulon that normally occurs during aging is necessary for lifespan extension by CR and overexpression of Ssd1.

**Figure 4.**
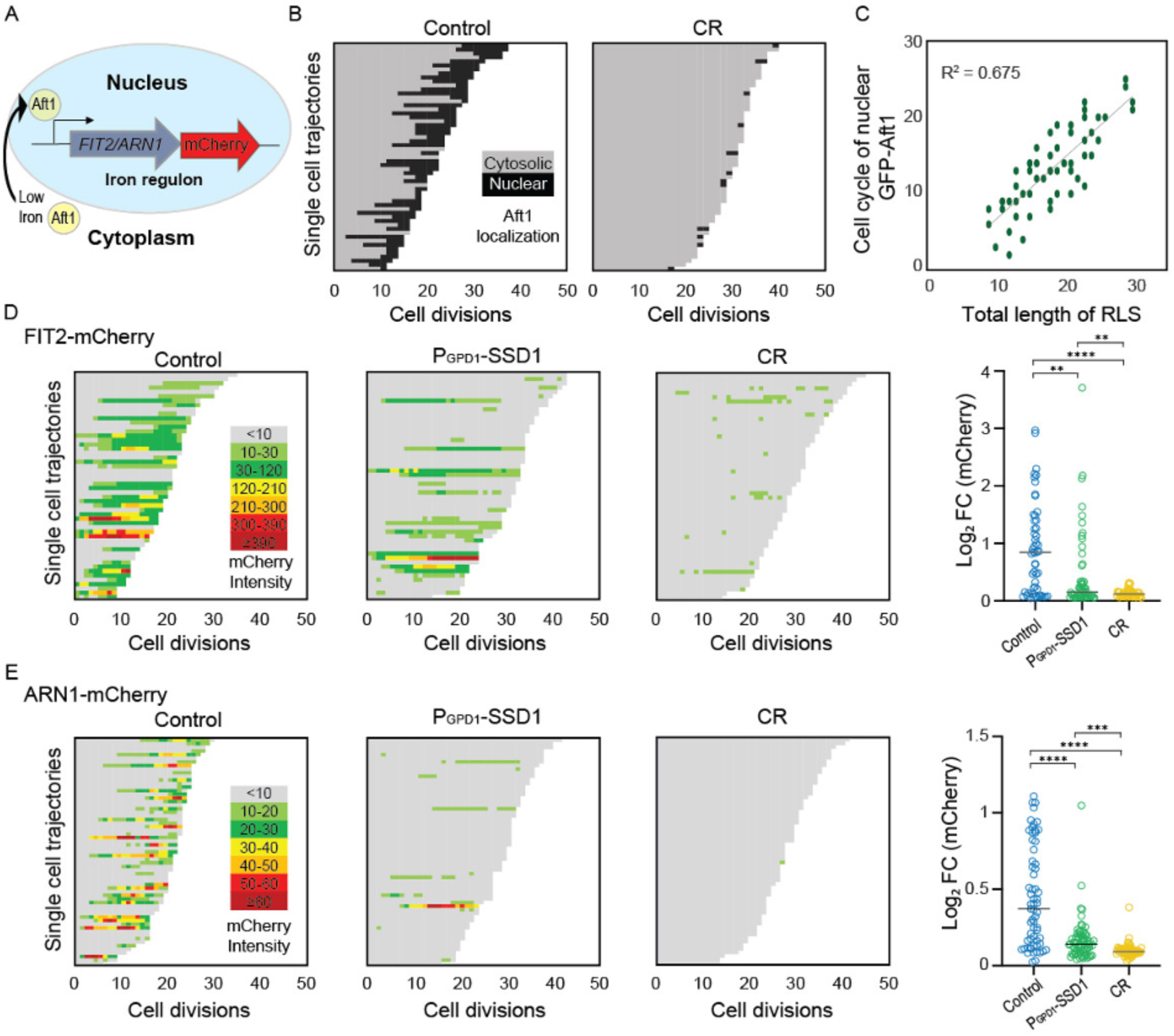
Age-dependent induction of the iron regulon predicts the end of lifespan, and is blocked by Ssd1 overexpression and CR. **A.** Schematic of iron regulon activation. **B** Single cell trajectories of nuclear localization of GFP-Aft1 during the RLS of control or CR yeast. **C.** Earlier nuclear localization of Aft1 in the lifespan correlates with a shorter total lifespan. Analysis of the data shown in panel B. **D.** Single cell trajectories of FIT2-mCherry expression during aging and Peak log fold change (FC) in mcherry intensity throughout the lifespan. **E.** Single cell trajectories of ARN1-mCherry expression during aging and fold change intensity throughout the lifespan. Statistical difference is indicated where n.s. is not significant change with p > 0.05, * p < 0.05, ** p ≤ 0.01, *** p ≤ 0.001 and **** p ≤ 0.0001.

**Figure 5.**
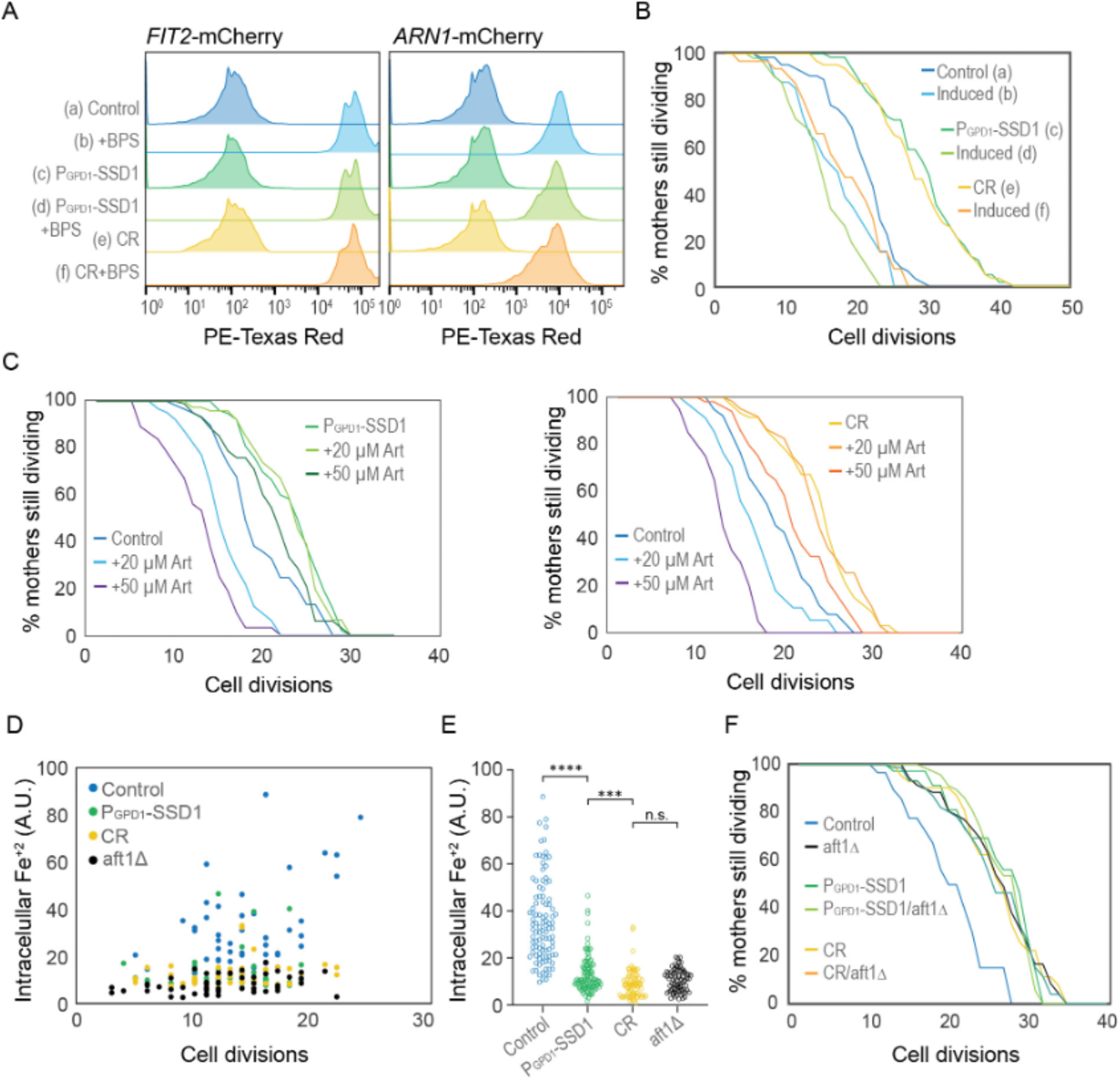
CR and Ssd1 overexpression extend lifespan by preventing the accumulation of iron resulting from age-dependent activation of the iron regulon. **A.** Pre-induction of iron regulon by BPS treatment measured by mCherry-tagged expression of Fit2 and Arn1, for 10000 cells per condition. **B.** RLS of cells that had the iron regulon induced or uninduced from panel A measured by Fit2-mcherry expression, for 63 (a), 41 (b), 63 (c), 50 (d), 63 (e), and 31 (f) cells, p-values of control vs induced: 0.002, P_G_poi-SSD1 vs induced: 0 and CR vs induced: 0. **C.** RLS of yeast overexpressing Ssd1 as well as under CR are protected from sensitivity to Artemisinin. Left plot: for 62 control, 64 20 µM Art, 37 50 µM Art, 86 Pgpdi-SSD1, 80 Pgpdi-SSD1 + 20 µM Art and 107 Pgpdi-SSD1 + 50 µM Art cells, p-values are control vs 20 µM Art: 4 x10-6, control vs 50 µM Art: 0, Pgpoi-SSD1 vs + P_G_pdi-SSD1 20 µM Art: 0.98 and Pgpoi-SSDI vs + P_G_pdi-SSD1 + 50 µM Art: 0.01. Right plot: 38 control, 34 20 µM Art, 30 50 µM Art, 61 CR, 61 CR + 20 µM Art, and 50 CR + 50 µM Art cells, p-values are control vs 20 µM Art: 0.01, control vs 50 µM Art: 1 x 10-7, CR vs CR 20 µM Art: 0.95 and CR vs + CR + 50 µM Art: 0.005. **D.** Relative levels of Fe^2+^ at indicated cell division of the RLS measured by Phen Green. **E.** Relative amount of Fe^2+^ in all cells of the indicated strains from panel D during the RLS, regardless of age. **F.** RLS of the indicated strains / conditions with and without aft1Δ, for 39 control, 71 aft1Δ, 37 P_G_pdi-SSD1, 72 P_G_pdi- SSD1/aft1Δ, 42 CR, and 74 CR/aft1Δ cells, p-values of control compared to all other strains are ≤ 1 x 10-4 while there is no significant difference between the strains / conditions other than the control.

### Intracellular iron accumulates during aging, but not in yeast overexpressing Ssd1 or during CR

Artemisinin (Art) is a malaria drug that becomes toxic upon reacting with intracellular iron^44^. We hypothesized that if control cells accumulated more iron during aging as a consequence of activation of the iron regulon, they would show higher sensitivity to Art than cells overexpressing Ssd1 or during CR which do not activate the iron regulon during aging to the same degree. Indeed, at 20 and 50 µM Art, control cells had a significantly reduced RLS compared to untreated controls (**Figure 5C**). In contrast, 20 µM Art had no effect on the RLS during CR or overexpressing Ssd1 and 50 µM Art only caused a modest decrease (**Figure 5C**). These results are consistent with CR and Ssd1 overexpression preventing the accumulation of free iron during aging that occurs when the iron regulon gets induced during aging.

To directly measure intracellular iron levels during aging, we used the fluorescent reporter Phen Green FL (PGFL), which identifies free labile iron (Fe^+2^)^33,45^. Control, P_GPD1_-SSD1, *aft1Δ* (a strain that is unable to upregulate cellular iron uptake because it is unable to activate the iron regulon), and CR-treated cells were aged in microfluidic chambers for ∼30 hours, generating a population ranging from 3 to ∼24 cell divisions in age. Cells were then treated with PGFL, followed by 1,10- phenanthroline^33^, an iron chelator. The difference in GFP fluorescence before and after PGFL treatment reflects the amount of free intracellular iron. We found that free labile iron accumulates during aging in control cells albeit with considerable cell-to-cell variability (**Figure 5D,E**). As a control, the *aft1Δ* cells did not accumulate free intracellular iron at any age (**Figure 5D,E**). This result indicates that the accumulation of free intracellular iron during aging is a consequence of activation of the iron regulon. By comparison to the control, the P_GPD1_-SSD1 and CR-treated cells consistently displayed lower intracellular iron concentrations than the control during aging (**Figure 5D,E**). Together these results indicate that there is an age-dependent iron intake due to activation of the iron regulon, which leads to a higher toxic intracellular free labile iron pool. Meanwhile, age-dependent activation of the iron regulon is reduced during CR or Ssd1 overexpression, limiting the uptake of iron during aging leading to beneficial effects on longevity.

### Failure to activate the iron regulon during aging is the molecular mechanism whereby Ssd1 overexpression and CR extend RLS

To determine whether reducing activation of the iron regulon is the only mechanism whereby CR and Ssd1 overexpression extend RLS, or whether additional mechanisms are at play, we performed epistasis analysis with the *aft1Δ* strain that is unable to induce the iron regulon. We deleted *AFT1* in both control and P_GPD1_-SSD1 strains and measured the RLS under both glucose rich and CR conditions. Given that *aft1Δ* strains have a growth defect independent from its role in iron regulon activation that is reversed by iron supplementation^27^, we included 200 µM FeCl^+3^ in our epistasis analyses. We found that *aft1Δ* strains grown in media supplemented with 200 µM FeCl^+3^ were long lived (**Figure 5F**), consistent with our prediction that failure to induce the iron regulon during aging is sufficient to extend lifespan. Importantly, combining *aft1Δ* with P_GPD1_-SSD1 led to the same degree of RLS extension as either *aft1Δ* or P_GPD1_-SSD1 alone (**Figure 5F**). Similarly, growing *aft1Δ* cells under CR led to the same degree of RLS extension as either *aft1Δ* or CR alone (**Figure 5F**). These epistasis analysis results indicate that the absence of activation of the iron regulon during aging, or muted iron regulon activation is the mechanism whereby CR and Ssd1 overexpression, respectively, extend the RLS (**Figure 6**).

**Figure 6.**
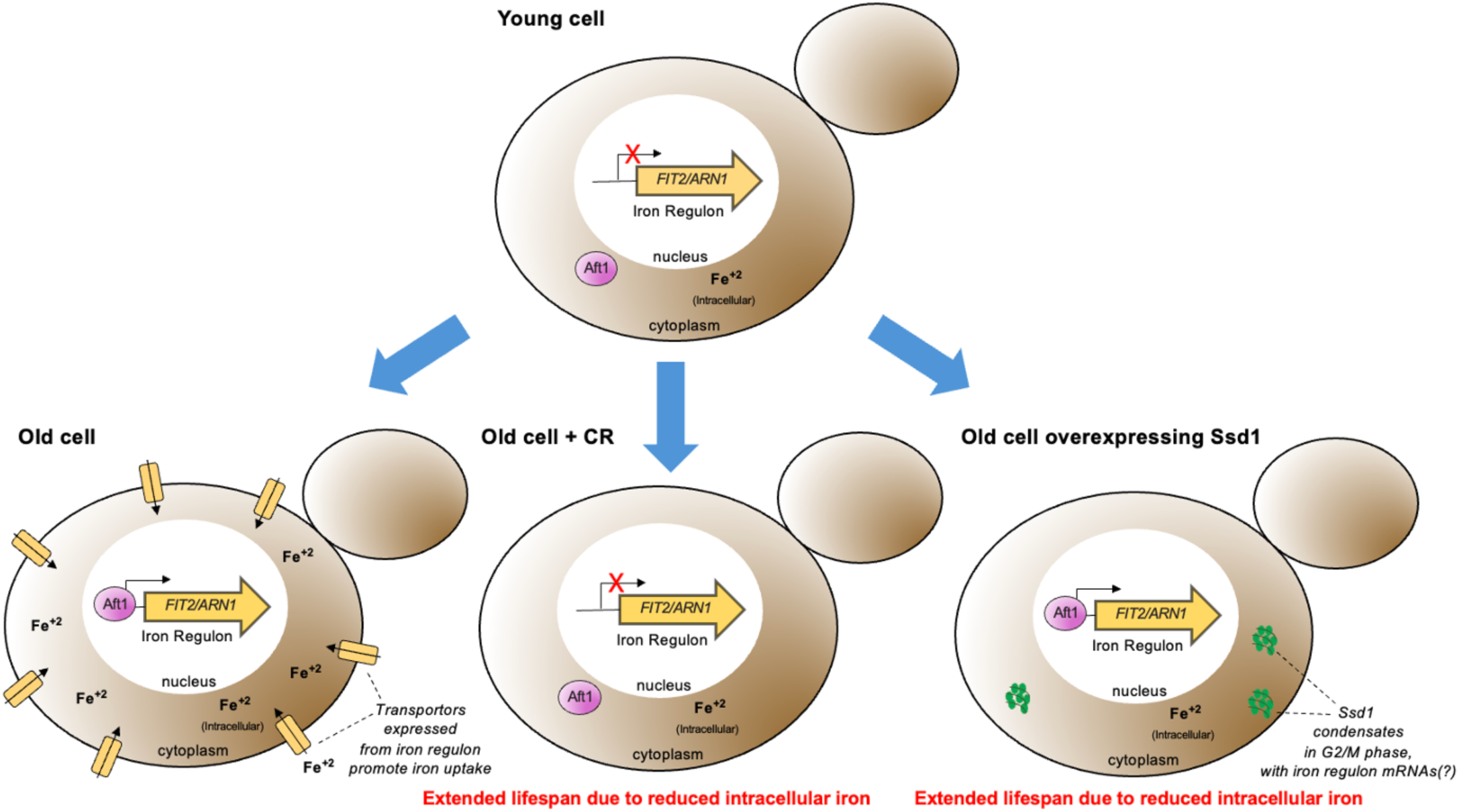
Model for molecular mechanism of RLS extension by Ssd1 overexpression and CR. During normal aging, cells have limited intracellular iron, potentially due to mitochondrial dysfunction reducing the assembly of iron-sulfur clusters, which leads to nuclear translocation of Aft1 and activation of the iron regulon. This subsequent deleterious increase in intracellular iron limits lifespan. CR also extends lifespan in a manner dependent on failure to activate the iron regulon; mechanistically CR indirectly prevents nuclear localization of Aft1, preventing production of the mRNAs from the iron regulon and the subsequent age-dependent increase in deleterious iron accumulation. In aging cells overexpressing Ssd1, transient cytoplasmic Ssd1 condensates appear that correlate with lifespan extension, where the lifespan extension is due to failure to activate the iron regulon preventing deleterious iron accumulation. We speculate that Ssd1 dephosphorylation by Sit4 after G phase likely leads to formation of Ssd1 condensates in aging cells overexpressing Ssd1 including mRNAs of the iron regulon genes *FIT2 and ARN1,* while dissolution of Ssd1 condensates occurs prior to cytokinesis upon Cbk1 -mediated phosphorylation of Ssd1.

## Discussion

Molecular insights into how anti-aging interventions achieve lifespan extension in any organism are scarce. Here we report that yeast replicative aging is accompanied by the accumulation of free labile iron, due to activation of the iron regulon – an event that foretells the end of lifespan. We reveal that CR and overexpression of the yeast mRNA binding protein Ssd1 both extend the yeast replicative lifespan by preventing activation of the iron regulon during aging, in turn preventing deleterious age-dependent accumulation of free labile iron that limits lifespan.

### Age-induced cell cycle-regulated Ssd1 condensates and lifespan extension

Lifespan extension by Ssd1 overexpression resulted in the age-induced transient appearance of Ssd1 cytoplasmic foci in a cell cycle stage-dependent manner (**Fig. 2, 3**). The fact that the Ssd1 foci could be dissolved with 1,6 Hexanediol and showed fusion properties is consistent with their being condensates. The cell cycle-dependent nature of formation and dissolution of the age-dependent Ssd1 condensates (**Fig. 2A**) is reminiscent of stress-induced formation of Ssd1 foci in cells overexpressing Ssd1 being triggered by Ssd1 dephosphorylation by the Sit4 phosphatase^34^. Sit4 functions in late G_1_ phase^46^ to promote a timely G_1_/S transition^47^, aligning nicely with the timing of appearance of age-induced Ssd1 foci after G_1_ phase (**Fig. 2A**). Moreover, stress-induced Ssd1 foci in cells overexpressing Ssd1 are dissolved by the Cbk1 kinase^10,34^, whose peak activity occurs immediately before cytokinesis^48^. This agrees with the fact that virtually all Ssd1 condensates that occurred in aging cells overexpressing Ssd1 dissolved by cytokinesis (**Fig. 2A**). It is tempting to speculate that not only is the cell cycle-dependent nature of the appearance of the Ssd1 foci beneficial for longevity, but perhaps also their dissolution by cytokinesis may be important, given that when Ssd1 condensates were retained through mitosis, it coincided with that being the last cell division of their lifespan (**Fig. 2A**).

The age-induced Ssd1 foci appear to be a novel type of condensate, because even though we could reproduce the colocalization of overexpressed Ssd1-GFP with stress-induced P-bodies and stress granules^10,34^, no such colocalization occurred during aging (**Fig. 2B-D**). This implies that the lifespan extension that occurs due to Ssd1 overexpression is likely mediated by the Ssd1 condensates themselves rather than by coaggregation with other stress-induced structures. It will be interesting to determine in the future exactly the nature of this novel type of Ssd1 condensate during aging and what kind of stress is triggering its formation during aging but not in young cells (**Fig. 2A**). Whatever the mechanism of their formation, age-dependent Ssd1 condensate formation is dependent on the overexpression of Ssd1, as it does not occur in cells expressing Ssd1-GFP from the endogenous promoter during aging, perhaps reflecting a disrupted balance between the molecular ratio of mRNAs and Ssd1 that may promote Ssd1 condensate formation. Notably, previous studies indicate that the fate of Ssd1-bound mRNAs depends on Ssd1’s aggregation state where the mRNAs bound to Ssd1 within a condensate are targeted for degradation or sequestration while the mRNAs bound to diffusive Ssd1 are targeted for translation^34,49^ consistent with our proposed mechanism whereby Ssd1 overexpression extends lifespan through beneficial targeted mRNA degradation / segregation to age-induced Ssd1 condensates (**Fig. 6**).

### Ssd1 overexpression and CR block the iron regulon, to extend lifespan

The induction of Fit2 and Arn1 that occurs during aging (**Fig. 4D,E**) was greatly reduced by overexpressing Ssd1 and CR (**Fig. 4D,E**) indicating that both anti-aging interventions are reducing activation of the iron regulon during aging. In agreement with induction of the iron regulon during aging, nuclear translocation of Aft1, one of the transcriptional activators of the iron regulon, occurs during aging and is a predictor of on average five remaining cell divisions in the lifespan (**Fig. 4B**). Moreover, the earlier the activation of the iron regulon in the lifespan, the shorter the lifespan (**Fig. 4C**). Notably, CR prevented nuclear translocation of Aft1 during aging (**Fig. 4B**) indicating that cells undergoing CR do not receive the signal to turn on the iron regulon during aging (**Fig. 6**). The signal for turning on the iron regulon during aging is likely the depletion of iron-sulfur clusters that occurs during aging as a consequence of mitochondrial dysfunction^26,50^. We speculate that the mechanism whereby CR prevents activation of the iron regulon is related to the fact that CR promotes mitochondrial biogenesis^51^ such that cells undergoing CR may not have depleted levels of iron-sulfur clusters during aging. CR releases the glucose-mediated repression of the heme activator protein (HAP) transcriptional activator complex required for transcription of key nuclear and mitochondrial genes needed for mitochondrial biogenesis^52^, while overexpression of the activating component of the HAP complex, Hap4, itself is sufficient to extend lifespan^52,53^ even in cells grown in glucose. It will be interesting to determine in the future whether overexpression of Hap4 also extends lifespan by preventing the induction of the iron regulon and deleterious accumulation of iron during aging. The ability of CR to activate mitophagy may also play a role in promoting mitochondrial biogenesis^54^.

While Ssd1 overexpression also reduces activation of the iron regulon to extend lifespan, we predict that a different mechanism is at play to block iron regulon activation than during CR. We speculate that the reason why Ssd1 overexpression reduces accumulation of Fit2 and Arn1 during aging (**Fig. 4D,E**) is that overexpressed Ssd1 may cause the degradation or sequestration of their mRNAs in the cytoplasmic age-induced Ssd1 condensates (**Fig. 6**). Fit2 and Arn1 are not among the known mRNAs that bind to Ssd1^11,12^ but this is to be expected as these previous studies were performed in conditions in which the iron regulon, and hence these transcripts, would not have been induced. Also, the Ssd1 mRNA binding partner studies had endogenous levels of Ssd1 not overexpressed which may bind to additional transcripts. Furthermore, it has been reported that Ssd1 can lead to the decay of mRNAs that it does not even bind to^49^ and this may be related to the reduced global protein synthesis that occurs due to Ssd1 overexpression^6^. Future studies are required to determine exactly how Ssd1 overexpression leads to reduced levels of Fit2 and Arn1 expression in aged cells.

Once the iron regulon is induced during aging, the production of high-affinity iron transporters such as Fit2 and Arn1 would lead to the subsequent rise in free labile intracellular iron during aging (**Fig. 5D,E**). Notably, free labile intracellular iron remains low during aging in strains with Ssd1 overexpression and under CR consistent with failure to activate the iron regulon (**Fig. 5D,E**). Together, these results suggest that age-related increases in iron uptake resulting from activation of the iron regulon are detrimental to replicative lifespan and that CR and Ssd1 overexpression extend lifespan by using different mechanisms to avoid activation of the iron regulon (**Fig. 6**). In agreement with their functioning through a shared pathway to extend lifespan, Ssd1 overexpression and CR are epistatic for lifespan extension (**Fig. 3**). The fact that pre-activating the iron regulon prevents lifespan extension by either CR or Ssd1 overexpression (**Fig. 5A,B**) is consistent with their preventing activation of the iron regulon being their sole mechanism of lifespan extension. Moreover, the key role of the iron regulon in limiting lifespan is apparent from the robust lifespan extension resulting from deletion of *AFT1*, which is also epistatic to the lifespan extension achieved by CR or Ssd1 overexpression (**Fig. 5F**).

### Iron levels and lifespan regulation

Activation of the iron regulon during aging has been proposed previously to shorten lifespan by repressing mitochondrial respiration via Cth2, a protein product of the iron regulon targeting the mRNAs of non-essential iron binding proteins for degradation^27^. However, we found that deletion of *CTH2* does not extend RLS to the same extent as overexpression of Ssd1 or CR (**Fig. 4-figure supplement 4**). Independent studies report that besides deletion of the gene encoding the iron regulon transcription factor *AFT1*^43^, single deletion of the *FIT2* or *ARN1* genes encoding high affinity iron transporters also extend lifespan^41,55,43^. Although *Saccharomyces cerevisiae* lacks iron export mechanisms, cytosolic iron can be sequestered into storage or iron-sulfur clusters^24^. It is important to note that the iron regulon is composed of several genes with, in some cases, opposite functions, for instance *CCC1*^56^ promotes vacuolar storage of iron, while *SMF3* promotes mobilization out of the vacuole^57^, coherently Ssd1 binds to *SMF3* mRNA^12^. As such, overexpressing Ssd1 *may* be modulating iron homeostasis beyond iron intake. Our data reveal that the amount of free labile iron tends to increase as yeast age whereas the level of free labile iron in cells lacking *ATF1* is low (**Fig. 5**), indicating that age-related iron regulon activation drives iron overload. We hypothesize that this excess free labile iron generates oxidative stress, damaging mitochondrial proteins, lipids, and DNA, ultimately impairing respiration. This mechanism is relevant to humans, as age-related iron accumulation in tissues, including the brain, is linked to cognitive decline^58^ and neurological disorders^59,60^, underscoring its importance for healthspan.

### Effects of CR on human iron metabolism

CR robustly extends lifespan from yeast to mammals^61^. While it is yet to be demonstrated that CR extends lifespan in humans, it is safe to say that promotes healthspan^13,62^. Several studies have shown age-related iron accumulation in human tissues, including the cerebral cortex^63^. Notably, recent studies demonstrate that CR reduces age-related iron accumulation in the brains of rhesus macaques^64^ and rodents^65,66^. In humans, CR and intermittent fasting improve memory and cognitive function during aging^67,68^. Whether these benefits stem from reduced iron accumulation remains to be investigated. We asked if there is any evidence that CR ameliorates toxic iron accumulation in humans. To investigate this, we analyzed published data. While we were unable to find proteomic data from specific tissues such as human brain under CR, we assessed the blood/plasma proteomics change of two studies from overweight donors subjected to CR leading to weight loss and stabilization^69,70^, two studies identifying proteins that correlate with age^71,72^ and one study analyzing protein signatures of centenarians^73^. Transferrin (TF), a blood plasma glycoprotein that binds and transports iron throughout the bloodstream to cells and tissues in the body, was upregulated in CR^69,70^, while decreases with age^71,72^, suggesting that an increase in TF may reflect enhanced capacity to bind and sequester free iron, reducing its toxic potential. CR also upregulates hepcidin (HAMP)^69^, a peptide hormone that serves as the master regulator of systemic iron homeostasis by inhibiting iron absorption and release from cells, consistent with people under CR potentially absorbing less iron from their diet^74^. Indeed, it has been shown that low levels of hepcidin leads to chronic liver disease as a result of iron accumulation^75^. HAMP was slightly but not significantly downregulated with age in both studies^71,72^. On the other hand, the transferrin receptor (TFRC) was downregulated in people under CR^69,70^, TFRC is present on cell surfaces to mediate iron uptake through transferrin-iron complexes^76^. A decrease under CR suggests that human cells reduce iron uptake from the plasma, limiting excess iron storage. TFRC is downregulated in centenarians^73^, and is slightly, but not significantly, increased with age^71^. Downregulation of ferritin heavy chain (FTH1) occurs in CR^69,70^, given that FTH1 is a component of ferritin, the iron storage protein, this downregulation could be a consequence of putative lower iron abundance in the plasma of people under CR. These findings highlight the potential for targeting iron related proteins to influence iron homeostasis and lifespan in humans. However, further studies will be needed to determine whether modulating the expression of these genes or proteins in humans could help reduce age-related iron accumulation in tissues, offering potential therapeutic strategies for conditions associated with iron dysregulation.

## Acknowledgements

We are indebted to Cornelia Kurischko for extensive thoughtful discussions about Ssd1 condensates, and the Weill Cornell Flow Cytometry Core for technical support. JKT is supported by NIH grants R01 AG079883 and R35 GM139816.

## Author contributions

JIG performed devised many of the experiments, performed the vast majority of the experiments with some assistance from CE, and wrote the first draft of the manuscript. JIG and JKT were responsible for program conception, support in data analysis and interpretation. JKT contributed to subsequent versions of the manuscript.

---

**Figure 1-figure supplement 1.**
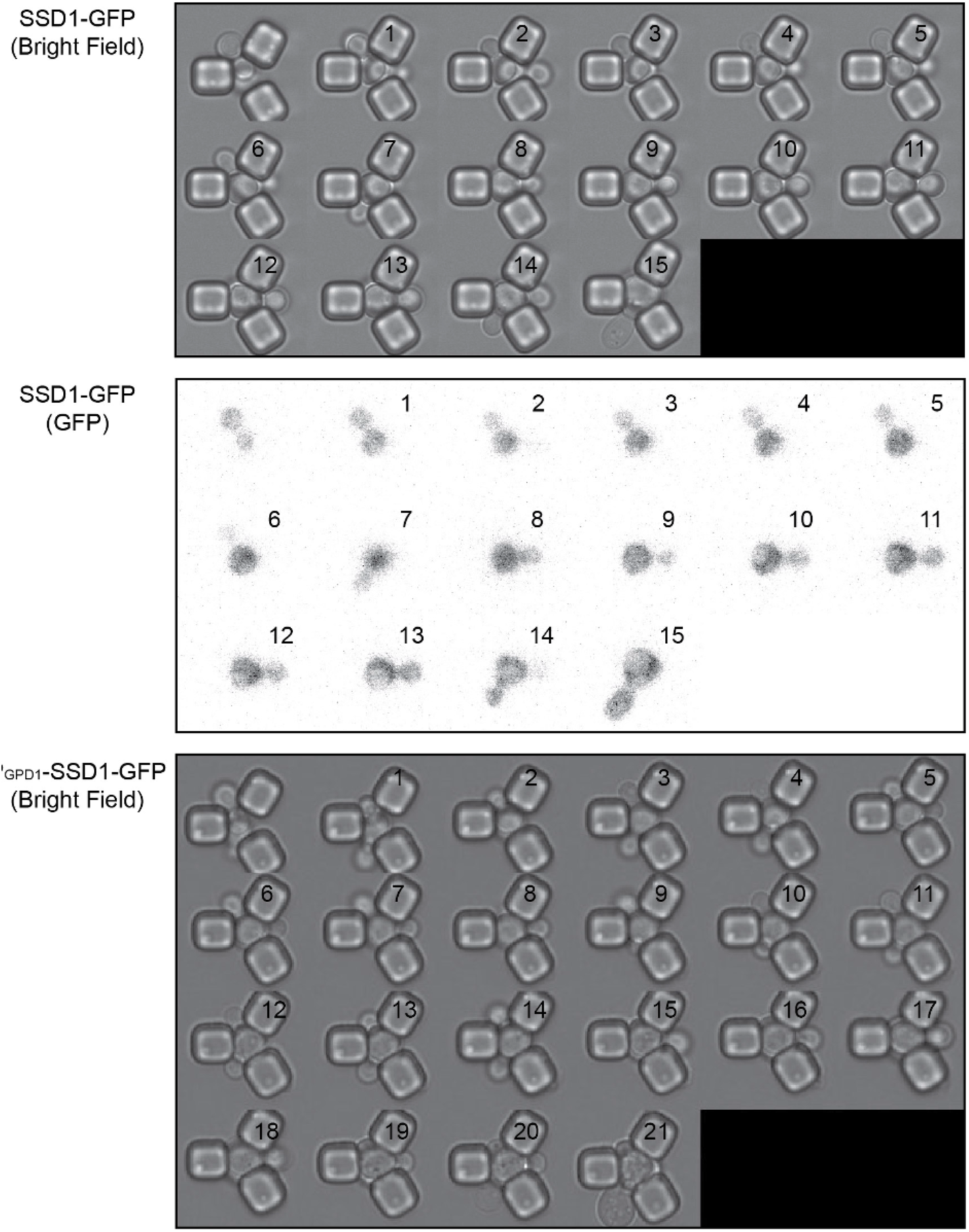
Single cell replicative aging of one cell, where images were taken for every cell cycle in its lifespan in the bright field and GFP channel for SSD1-GFP and in the bright field for P_GPD1_- SSD1-GFP and corresponds to image taken in the GFP channel shown in Fig-1C.

**Figure 1-figure supplement 2.**
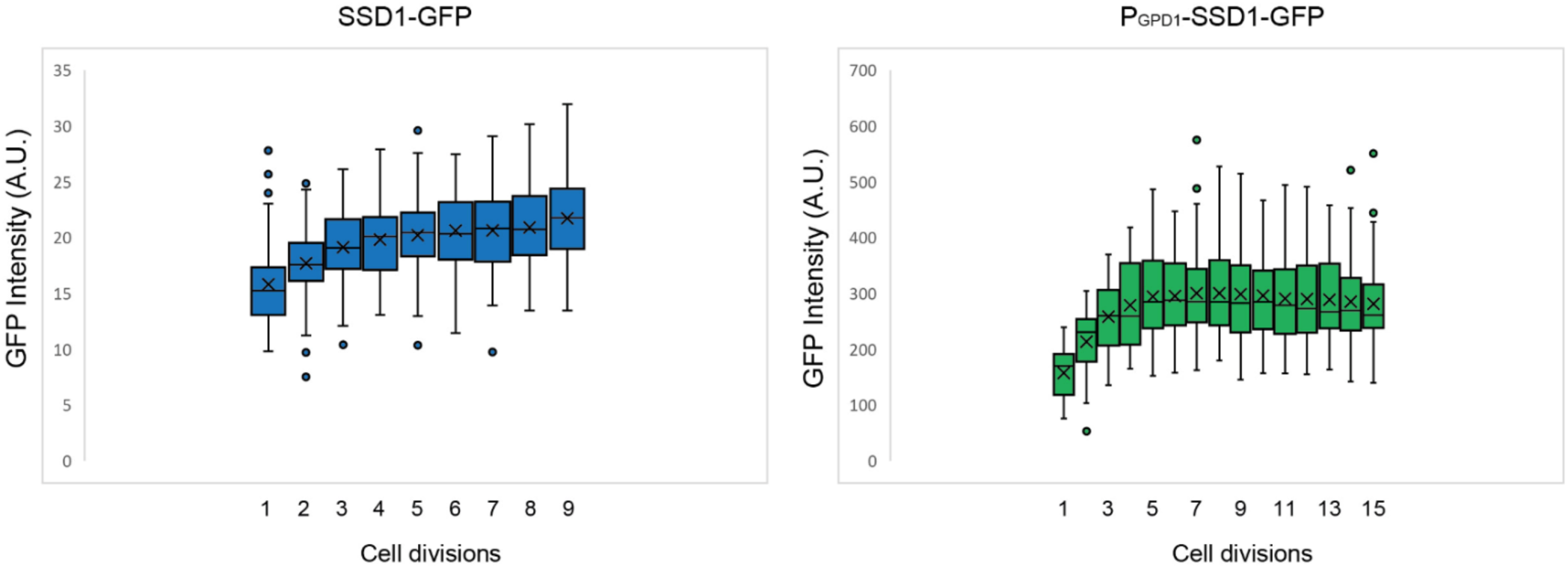
Relative SSD1-GFP protein amount per cell throughout the lifespan, based on total cellular fluorescence intensity of strains SSD1-GFP and P_G_pdi-SSD1-GFP during the cell division in which 100% of the population (mothers) are actively dividing for the data shown in Fig. 1D. Average and standard deviation for 63 cells per strain is shown. The box indicates interquartile range (IQR), whiskers extend to the smallest and largest non-outlier values and outliers are plotted as individual points outside Quartile(Q)1-1.5×IQR or Q3+1.5×IQR, where IQR = Q3-Q1. Data are shown for cell divisions in which 100% of the mothers are still dividing.

**Figure 1-figure supplement 3.**
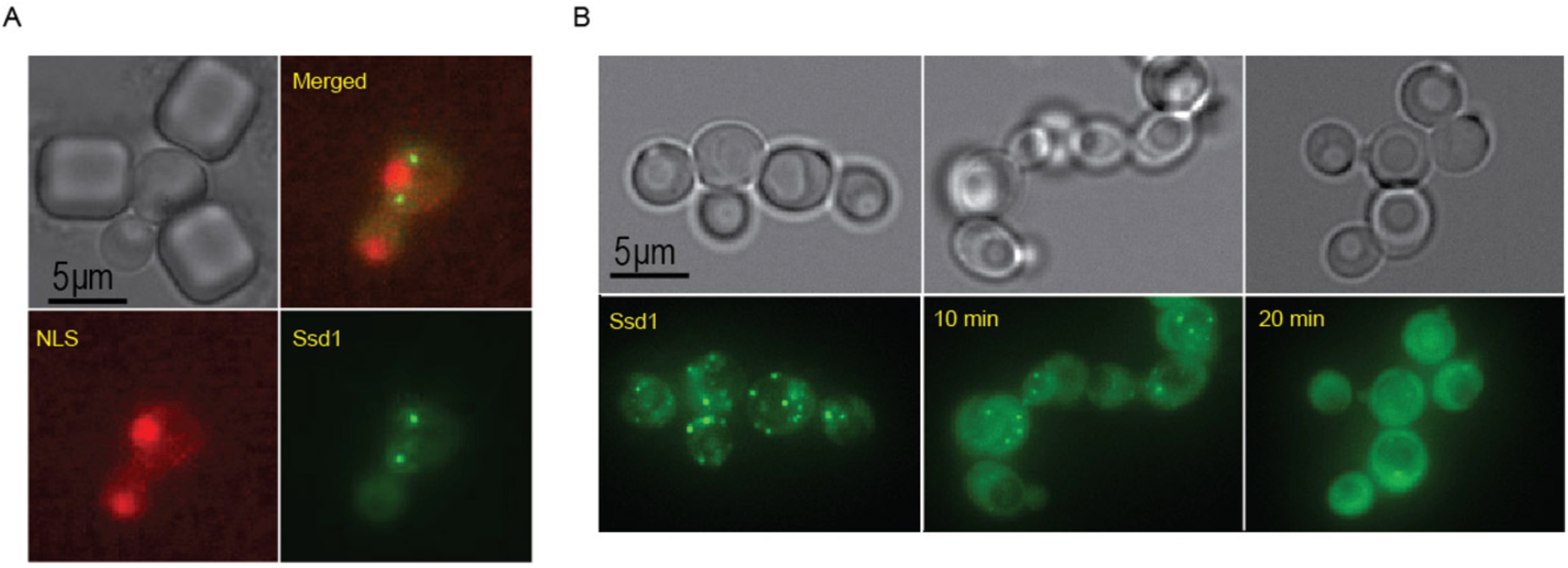
**A.** Ssd1-GFP foci formed during aging are cytoplasmic. mCherry-NLS indicates the nucleus. **B.** 1-6hexanediol dissolves Ssd1-GFP foci, images taken before, or at 10 or 20 mins after, addition of 1-6hexanediol.

**Figure 2-figure supplement 1.**
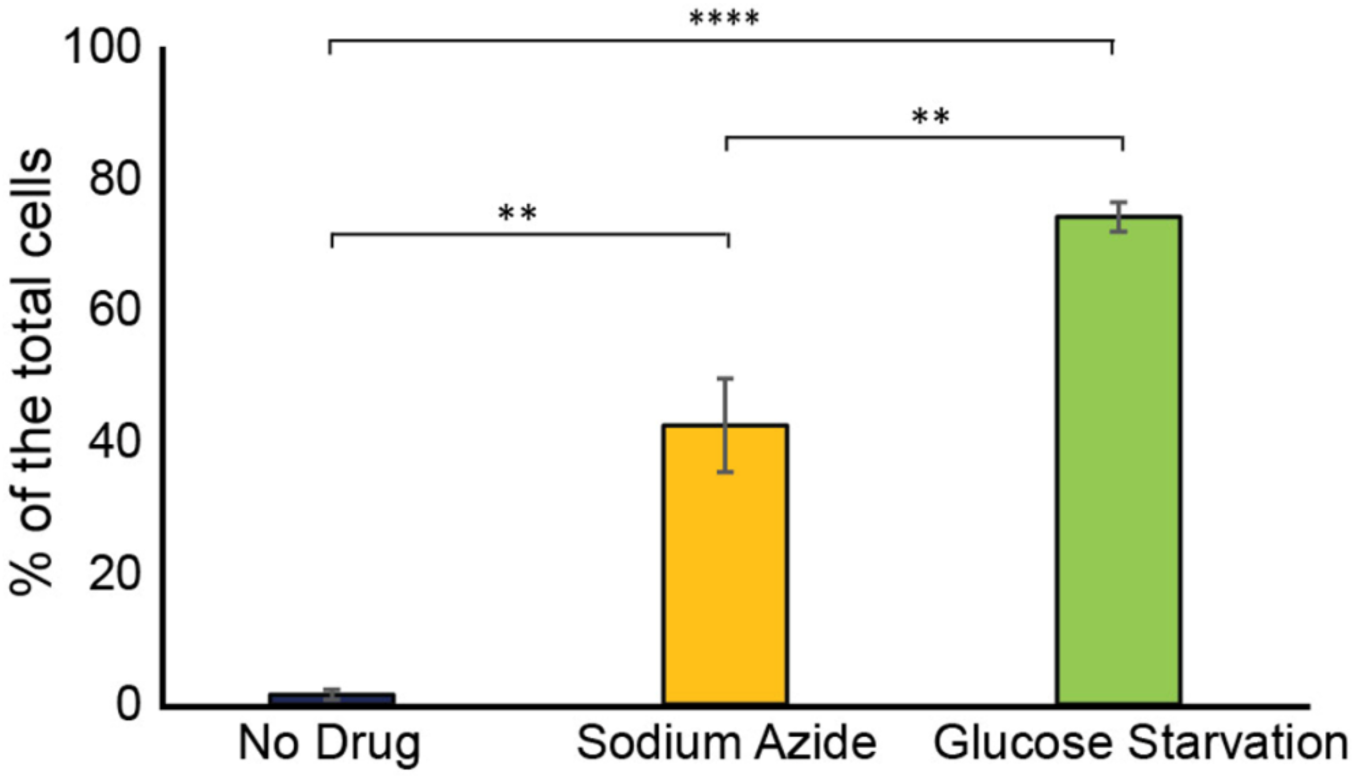
Frequency of Ssd1 foci formation in exponentially growing cells, treated with sodium azide and under glucose starvation in strain P_gpd1_-SSD1-GFP. Average and standard deviation are shown for analysis of biological triplicate experiments for >60 cells per condition per experiment, p-values determined by Student’s t-test are indicated by ** for 0.01 and ***** for 0.0001.

**Figure 4-figure supplement 1.**
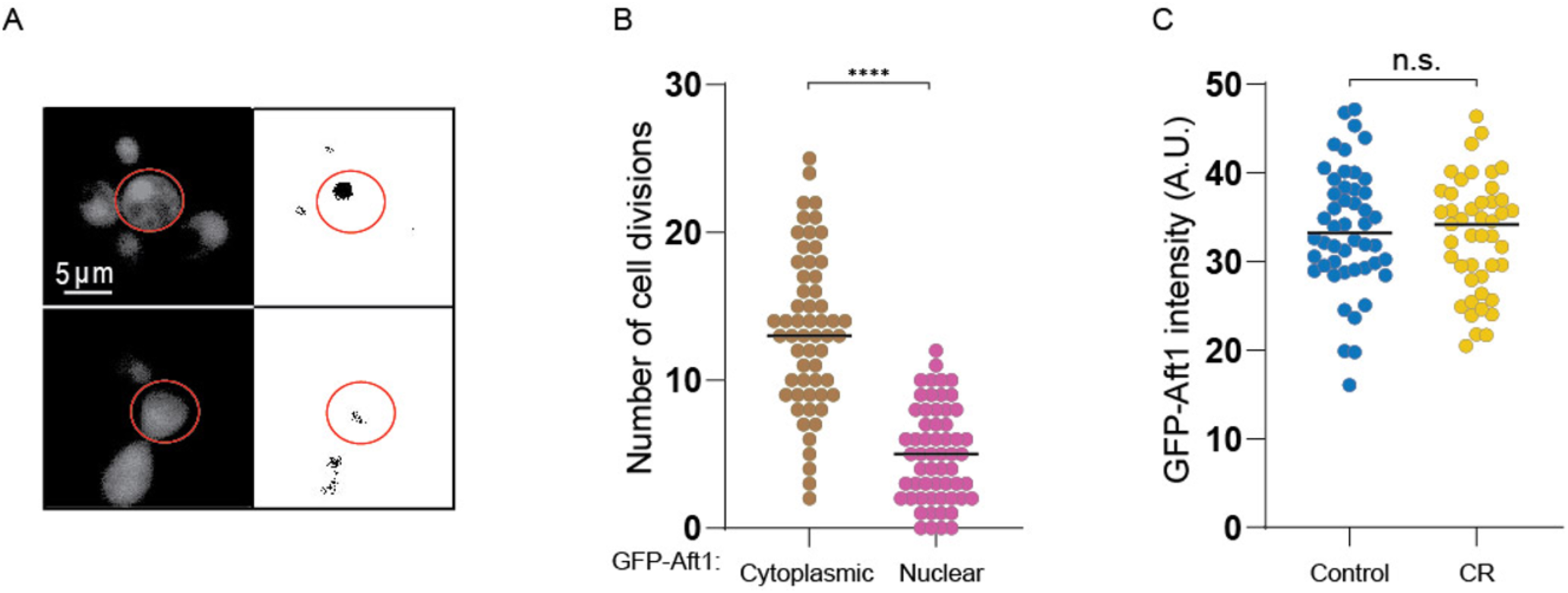
**A.** Example of old cell with (top) and without (bottom) nuclear localization of GFP-Aft1 on the left, profile after threshold (see materials and methods) on the right. **B.** Localization of Aft1 to the nucleus occurs on average 5 divisions before the end of lifespan. The analysis is of the data shown in Fig. 4B. **C.** CR does not affect GFP-Aft1 expression as determined by fluorescence intensity. 50 cells were analyzed per conditions in each plot. n.s. indicates a not significant change of p > 0.05 and **** is p ≤ 0.0001.

**Figure 4-figure supplement 2.**
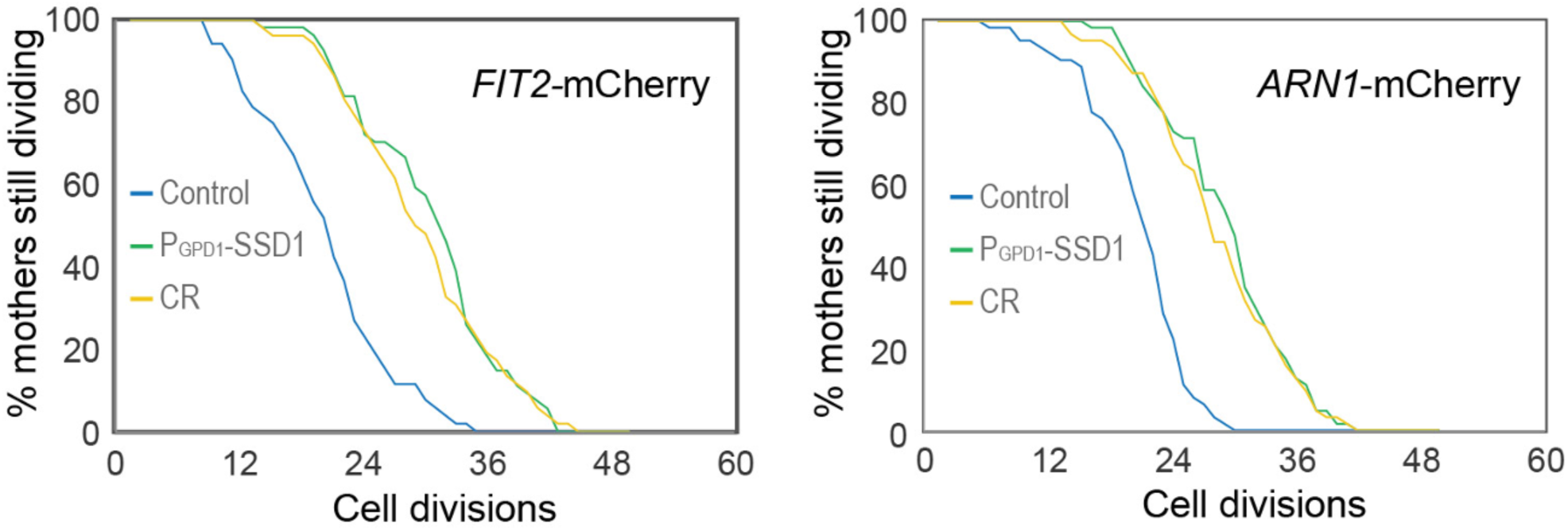
Replicative lifespan for experiments shown in Figure 4D and 4E. p-values for both graphs of control versus P_gpd_-SSD1 and CR is 0, while for P_gpd_-SSD1 versus CR is not significant, 50 cells were analyzed for each condition.

**Figure 4-figure supplement 3.**
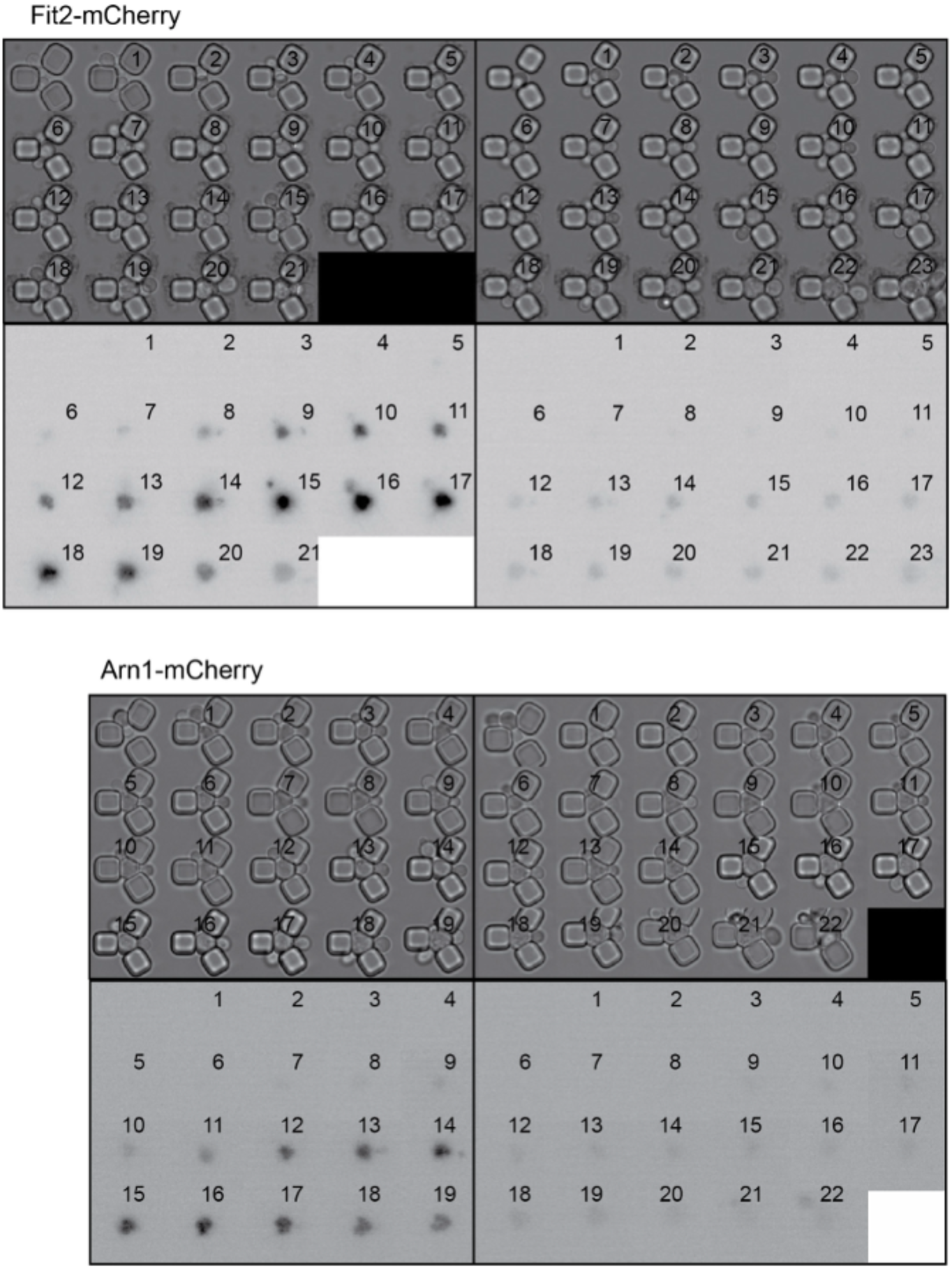
Single cell examples of experiments shown in Figure 4D and 4E, to show variability between different cells. The number indicates the cell division number in the RLS. Top is imaged under bright light and the bottom under mCherry fluorescence detection.

**Figure 4-figure supplement 4.**
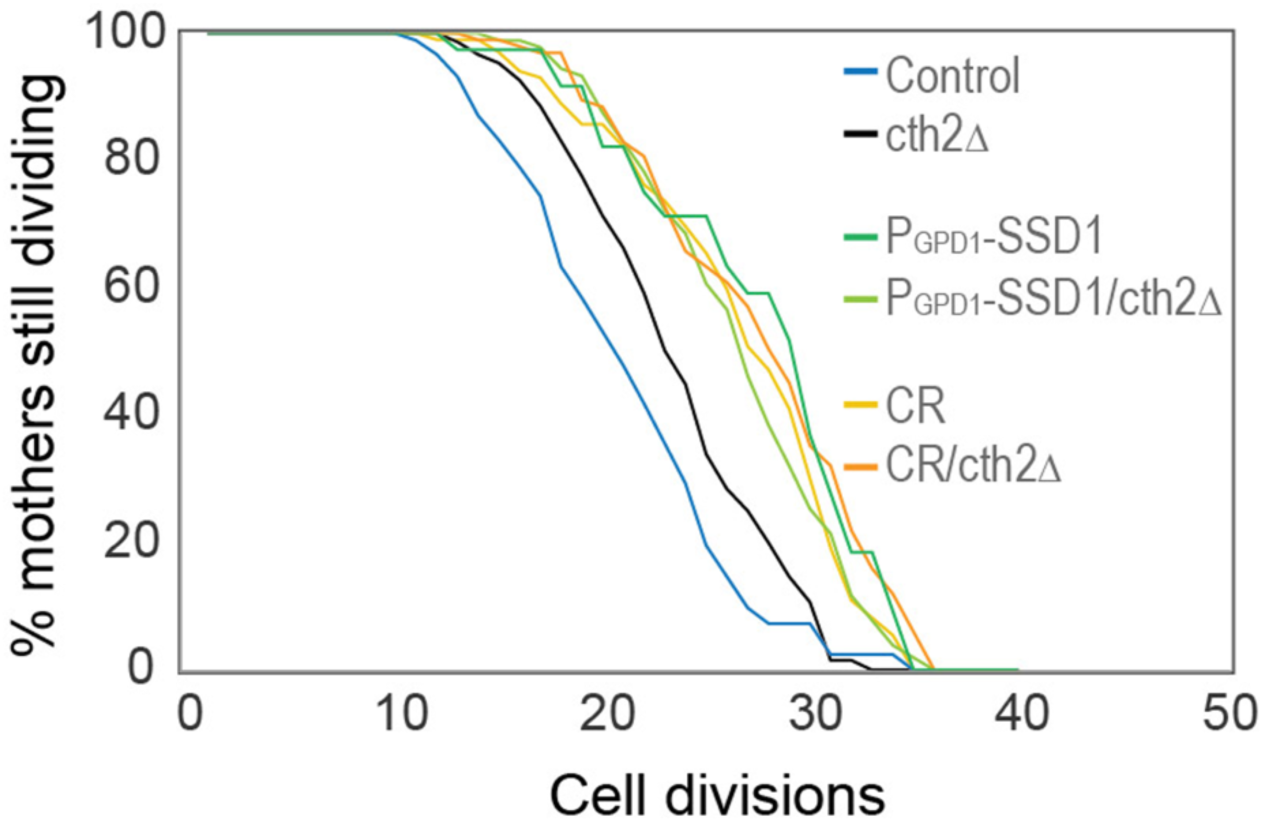
RLS of the indicated strains / conditions with and without *cth2Δ,* the number of cells counted per condition are control: 91, cth2Δ: 150, P_gpd1_-SSD1: 39, P_GPD1_- SSD1/cth2Δ: 91, CR: 101, CR/cth2Δ: 98, p-values: control compared to cth2Δ is 6 x 10^-3^, *cth2Δ* compared to CR, P_GPD1_- SSD1/cth2Δ, cth2Δ under CR and P_gpd1_-SSD1 are 5 x 10^-6^, 1×10^-5^, 3 x 10^-8^ and 2 x 10^-4^, respectively.

**Supplemental Movie 1.** Bright field (**A**.) and corresponding fluorescence microscopy (**B**.) of Ssd1-GFP for an SSD1-GFP cell during the RLS. Bright field (**C**.) and corresponding fluorescence microscopy (**D**.) of Ssd1-GFP for a P GPD1 -SSD1-GFP cell during the RLS. Numbers indicate the number of cell divisions into the lifespan.

**Supplemental Movie 2.** Fluorescence microscopy of Ssd1-GFP condensates undergoing fusion in an aging PGPD1-SSD1-GFP cell.

**Supplemental Table 1:** Yeast strains used in this study.

|  |  |
| --- | --- |
| IGY138 | MATα *his3Δ leu2Δ ura3Δ ssd1::NatMX* |
| IGY081 | MATα *his3Δ leu2Δ ura3Δ SSD1-GFP (KanMX6)* |
| IGY086 | MATα *his3Δ leu2Δ ura3Δ P<sub>GPD1</sub>-SSD1-GFP (KanMX6)* |
| IGY536 | MATα *his3Δ leu2Δ ura3Δ SSD1-mCherry (HIS3)* |
| IGY540 | MATα *his3Δ leu2Δ ura3Δ P<sub>GPD1</sub>-SSD1-mCherry (HIS3)* |
| IGY106 | MATα *his3Δ leu2Δ ura3Δ SSD1-GFP (KanMX6) PAB1-mCherry (HIS3)* |
| IGY115 | MATα *his3Δ leu2Δ ura3Δ P<sub>GPD1</sub>-SSD1-GFP (KanMX6) PAB1-mCherry (HIS3)* |
| IGY107 | MATα *his3Δ leu2Δ ura3Δ SSD1-GFP (KanMX6) EDC3-mCherry (HIS3)* |
| IGY110 | MATα *his3Δ leu2Δ ura3Δ P<sub>GPD1</sub>-SSD1-GFP (KanMX6) EDC3-mCherry (HIS3)* |
| IGY117 | MATα *his3Δ leu2Δ ura3Δ SSD1-GFP (KanMX6) HSP104-mCherry (HIS3)* |
| IGY111 | MATα *his3Δ leu2Δ ura3Δ P<sub>GPD1</sub>-SSD1-GFP (KanMX6) HSP104-mCherry (HIS3)* |
| IGY544 | MATα *his3Δ leu2Δ ura3Δ SSD1-mCherry (HIS3) URA3-P<sub>GPD1</sub>-GFP_AFT1* |
| IGY545 | MATα *his3Δ leu2Δ ura3Δ P<sub>GPD1</sub>-SSD1-GFP-mCherry (HIS3) URA3-P<sub>GPD1</sub>-GFP_AFT1* |
| IGY521 | MATα *his3Δ leu2Δ ura3Δ SSD1-GFP (KanMX6) ARN1-mCherry (HIS3)* |
| IGY522 | MATα *his3Δ leu2Δ ura3Δ P<sub>GPD1</sub>-SSD1-GFP (KanMX6) ARN1-mCherry (HIS3)* |
| IGY529 | MATα *his3Δ leu2Δ ura3Δ SSD1-GFP (KanMX6) FIT2-mCherry (HIS3)* |
| IGY530 | MATα *his3Δ leu2Δ ura3Δ P<sub>GPD1</sub>-SSD1-GFP (KanMX6) FIT2-mCherry (HIS3)* |
| IGY516 | MATα *his3Δ leu2Δ ura3Δ SSD1-GFP (KanMX6) aft1::HIS3* |
| IGY524 | MATα *his3Δhis3Δ leu2Δ ura3Δ* P<sub>GPD1</sub>-SSD1-GFP (KanMX6)<br>*aft1::HIS3* |

## Reference List

1 Anderson, R. M. & Weindruch, R. Metabolic reprogramming, caloric restriction and aging. Trends Endocrinol Metab 21, 134–141 (2010). 10.1016/j.tem.2009.11.005

2 Blagosklonny, M. V. Rapamycin for longevity: opinion article. Aging (Albany NY) 11, 8048–8067 (2019). 10.18632/aging.102355

3 Lopez-Otin, C., Blasco, M. A., Partridge, L., Serrano, M. & Kroemer, G. Hallmarks of aging: An expanding universe. Cell 186, 243–278 (2023). 10.1016/j.cell.2022.11.001

4 López-Otín, C., Blasco, M. A., Partridge, L., Serrano, M. & Kroemer, G. in Cell Vol. 153 1194–1194 (Elsevier B.V., 2013).

5 Anisimova, A. S. et al. Multifaceted deregulation of gene expression and protein synthesis with age. Proc Natl Acad Sci U S A 117, 15581–15590 (2020). 10.1073/pnas.2001788117

6 Hu, Z. et al. Ssd1 and Gcn2 suppress global translation efficiency in replicatively aged yeast while their activation extends lifespan. eLife 7 (2018). 10.7554/eLife.35551

7 Luukkonen, B. G. & Seraphin, B. A conditional U5 snRNA mutation affecting pre-mRNA splicing and nuclear pre-mRNA retention identifies SSD1/SRK1 as a general splicing mutant suppressor. Nucleic Acids Res 27, 3455–3465 (1999). 10.1093/nar/27.17.3455

8 Kaeberlein, M., Andalis, A. A., Liszt, G. B., Fink, G. R. & Guarente, L. Saccharomyces cerevisiae SSD1-V Confers Longevity by a Sir2p-Independent Mechanism. Genetics 166, 1661–1672 (2004). 10.1093/genetics/166.4.1661

9 Steinkraus, K. A., Kaeberlein, M. & Kennedy, B. K. Replicative aging in yeast: the means to the end. Annu Rev Cell Dev Biol 24, 29–54 (2008). 10.1146/annurev.cellbio.23.090506.123509

10 Kurischko, C., Kim, H. K., Kuravi, V. K., Pratzka, J. & Luca, F. C. The yeast Cbk1 kinase regulates mRNA localization via the mRNA-binding protein Ssd1. J Cell Biol 192, 583–598 (2011). 10.1083/jcb.201011061

11 Hogan, D. J., Riordan, D. P., Gerber, A. P., Herschlag, D. & Brown, P. O. Diverse RNA-binding proteins interact with functionally related sets of RNAs, suggesting an extensive regulatory system. PLoS Biol 6, e255 (2008). 10.1371/journal.pbio.0060255

12 Dutcher, H. A. & Gasch, A. P. Investigating the role of RNA-binding protein Ssd1 in aneuploidy tolerance through network analysis. RNA 31, 100–112 (2024). 10.1261/rna.080199.124

13 Caristia, S. et al. Is Caloric Restriction Associated with Better Healthy Aging Outcomes? A Systematic Review and Meta-Analysis of Randomized Controlled Trials. Nutrients 12 (2020). 10.3390/nu12082290

14 Colman, R. J. et al. Caloric restriction reduces age-related and all-cause mortality in rhesus monkeys. Nat Commun 5, 3557 (2014). 10.1038/ncomms4557

15 Ungvari, Z., Parrado-Fernandez, C., Csiszar, A. & de Cabo, R. Mechanisms underlying caloric restriction and lifespan regulation: implications for vascular aging. Circ Res 102, 519–528 (2008). 10.1161/CIRCRESAHA.107.168369

16 Mesquita, A. et al. Caloric restriction or catalase inactivation extends yeast chronological lifespan by inducing H2O2 and superoxide dismutase activity. Proc Natl Acad Sci U S A 107, 15123–15128 (2010). 10.1073/pnas.1004432107

17 Hughes, A. L. & Gottschling, D. E. An early age increase in vacuolar pH limits mitochondrial function and lifespan in yeast. Nature 492, 261–265 (2012). 10.1038/nature11654

18 Choi, J. S. & Lee, C. K. Maintenance of cellular ATP level by caloric restriction correlates chronological survival of budding yeast. Biochem Biophys Res Commun 439, 126–131 (2013). 10.1016/j.bbrc.2013.08.014

19 An, H. S. et al. Caloric restriction reverses left ventricular hypertrophy through the regulation of cardiac iron homeostasis in impaired leptin signaling mice. Sci Rep 10, 7176 (2020). 10.1038/s41598-020-64201-2

20 Ben Zichri-David, S., Shkuri, L. & Ast, T. Pulling back the mitochondria’s iron curtain. NPJ Metab Health Dis 3, 6 (2025). 10.1038/s44324-024-00045-y

21 Chutvanichkul, B., Vattanaviboon, P., Mas-Oodi, S., Y, U. P. & Wanachiwanawin, W. Labile iron pool as a parameter to monitor iron overload and oxidative stress status in beta-thalassemic erythrocytes. Cytometry B Clin Cytom 94, 631–636 (2018). 10.1002/cyto.b.21633

22 Martinez-Garay, C. A., de Llanos, R., Romero, A. M., Martinez-Pastor, M. T. & Puig, S. Responses of Saccharomyces cerevisiae Strains from Different Origins to Elevated Iron Concentrations. Appl Environ Microbiol 82, 1906–1916 (2016). 10.1128/AEM.03464-15

23 Yamaguchi-Iwai, Y., Ueta, R., Fukunaka, A. & Sasaki, R. Subcellular localization of Aft1 transcription factor responds to iron status in Saccharomyces cerevisiae. J Biol Chem 277, 18914–18918 (2002). 10.1074/jbc.M200949200

24 Ramos-Alonso, L., Romero, A. M., Martinez-Pastor, M. T. & Puig, S. Iron Regulatory Mechanisms in Saccharomyces cerevisiae. Front Microbiol 11, 582830 (2020). 10.3389/fmicb.2020.582830

25 Rutherford, J. C. et al. Activation of the iron regulon by the yeast Aft1/Aft2 transcription factors depends on mitochondrial but not cytosolic iron-sulfur protein biogenesis. J Biol Chem 280, 10135–10140 (2005). 10.1074/jbc.M413731200

26 Chen, K. L. et al. Loss of vacuolar acidity results in iron-sulfur cluster defects and divergent homeostatic responses during aging in Saccharomyces cerevisiae. Geroscience 42, 749–764 (2020). 10.1007/s11357-020-00159-3

27 Patnaik, P. K. et al. Deficiency of the RNA-binding protein Cth2 extends yeast replicative lifespan by alleviating its repressive effects on mitochondrial function. Cell Rep 40, 111113 (2022). 10.1016/j.celrep.2022.111113

28 Baker Brachmann, C., et al. Designer deletion strains derived from Saccharomyces cerevisiae S288C: A useful set of strains and plasmids for PCR-mediated gene disruption and other applications. Yeast 14, 115–132 (1998). 10.1002/(SICI)1097-0061(19980130)14:2<115::AID-YEA204>3.0.CO;2-2

29 Longtine, M. S. et al. Additional modules for versatile and economical PCR-based gene deletion and modification in Saccharomyces cerevisiae. Yeast 14, 953–961 (1998). 10.1002/(SICI)1097-0061(199807)14:10<953::AID-YEA293>3.0.CO;2-U

30 Chee, M. K. & Haase, S. B. New and Redesigned pRS Plasmid Shuttle Vectors for Genetic Manipulation of Saccharomycescerevisiae. G3 (Bethesda) 2, 515–526 (2012). 10.1534/g3.111.001917

31 Gutierrez, J. I. & Tyler, J. K. A mortality timer based on nucleolar size triggers nucleolar integrity loss and catastrophic genomic instability. Nature Aging 4, 1782–1793 (2024). 10.1038/s43587-024-00754-5

32 Han, S. K. et al. OASIS 2: online application for survival analysis 2 with features for the analysis of maximal lifespan and healthspan in aging research. Oncotarget 7, 56147–56152 (2016). 10.18632/oncotarget.11269

33 Gomez-Gallardo, M., Sanchez, L. A., Diaz-Perez, A. L., Cortes-Rojo, C. & Campos-Garcia, J. Data on the role of iba57p in free Fe(2+) release and O(2)(·-) generation in Saccharomyces cerevisiae. Data Brief 18, 198–202 (2018). 10.1016/j.dib.2018.03.023

34 Kurischko, C. & Broach, J. R. Phosphorylation and nuclear transit modulate the balance between normal function and terminal aggregation of the yeast RNA-binding protein Ssd1. Mol Biol Cell 28, 3057–3069 (2017). 10.1091/mbc.E17-02-0100

35 Hose, J. et al. The genetic basis of aneuploidy tolerance in wild yeast. Elife 9 (2020). 10.7554/eLife.52063

36 Saarikangas, J. & Barral, Y. Protein aggregates are associated with replicative aging without compromising protein quality control. Elife 4 (2015). 10.7554/eLife.06197

37 Prasad, T., Chandra, A., Mukhopadhyay, C. K. & Prasad, R. Unexpected link between iron and drug resistance of Candida spp.: iron depletion enhances membrane fluidity and drug diffusion, leading to drug-susceptible cells. Antimicrob Agents Chemother 50, 3597–3606 (2006). 10.1128/AAC.00653-06

38 Ueta, R., Fukunaka, A. & Yamaguchi-Iwai, Y. Pse1p mediates the nuclear import of the iron-responsive transcription factor Aft1p in Saccharomyces cerevisiae. J Biol Chem 278, 50120–50127 (2003). 10.1074/jbc.M305046200

39 Diab, H. I. & Kane, P. M. Loss of vacuolar H+-ATPase (V-ATPase) activity in yeast generates an iron deprivation signal that is moderated by induction of the peroxiredoxin TSA2. J Biol Chem 288, 11366–11377 (2013). 10.1074/jbc.M112.419259

40 Gil, F. N., Belli, G. & Viegas, C. A. The Saccharomyces cerevisiae response to stress caused by the herbicidal active substance alachlor requires the iron regulon transcription factor Aft1p. Environ Microbiol 19, 485–499 (2017). 10.1111/1462-2920.13439

41 McCormick, M. A. et al. A Comprehensive Analysis of Replicative Lifespan in 4,698 Single-Gene Deletion Strains Uncovers Conserved Mechanisms of Aging. Cell Metab 22, 895–906 (2015). 10.1016/j.cmet.2015.09.008

42 Olmez, T. T. et al. Sis2 regulates yeast replicative lifespan in a dose-dependent manner. Nat Commun 14, 7719 (2023). 10.1038/s41467-023-43233-y

43 Yu, R. et al. Inactivating histone deacetylase HDA promotes longevity by mobilizing trehalose metabolism. Nature Communications 12, 1981–1981 (2021). 10.1038/s41467-021-22257-2

44 Chen, G. Q. et al. Artemisinin compounds sensitize cancer cells to ferroptosis by regulating iron homeostasis. Cell Death Differ 27, 242–254 (2020). 10.1038/s41418-019-0352-3

45 Jing, X. et al. HIF-2alpha/TFR1 mediated iron homeostasis disruption aggravates cartilage endplate degeneration through ferroptotic damage and mtDNA release: A new mechanism of intervertebral disc degeneration. J Orthop Translat 46, 65–78 (2024). 10.1016/j.jot.2024.03.005

46 Sutton, A., Immanuel, D. & Arndt, K. T. The SIT4 protein phosphatase functions in late G1 for progression into S phase. Molecular and Cellular Biology 11, 2133–2148 (1991). 10.1128/mcb.11.4.2133

47 Jiang, Y. Regulation of the cell cycle by protein phosphatase 2A in Saccharomyces cerevisiae. Microbiol Mol Biol Rev 70, 440–449 (2006). 10.1128/MMBR.00049-05

48 Brace, J., Hsu, J. & Weiss, E. L. Mitotic exit control of the Saccharomyces cerevisiae Ndr/LATS kinase Cbk1 regulates daughter cell separation after cytokinesis. Mol Cell Biol 31, 721–735 (2011). 10.1128/MCB.00403-10

49 Jansen, J. M., Wanless, A. G., Seidel, C. W. & Weiss, E. L. Cbk1 regulation of the RNA-binding protein Ssd1 integrates cell fate with translational control. Curr Biol 19, 2114–2120 (2009). 10.1016/j.cub.2009.10.071

50 Veatch, J. R., McMurray, M. A., Nelson, Z. W. & Gottschling, D. E. Mitochondrial Dysfunction Leads to Nuclear Genome Instability via an Iron-Sulfur Cluster Defect. Cell 137, 1247–1258 (2009). 10.1016/j.cell.2009.04.014

51 Lopez-Lluch, G. et al. Calorie restriction induces mitochondrial biogenesis and bioenergetic efficiency. Proc Natl Acad Sci U S A 103, 1768–1773 (2006). 10.1073/pnas.0510452103

52 Lin, S. J. et al. Calorie restriction extends Saccharomyces cerevisiae lifespan by increasing respiration. Nature 418, 344–348 (2002). 10.1038/nature00829

53 Li, Y. et al. A programmable fate decision landscape underlies single-cell aging in yeast. Science 369, 325–329 (2020). 10.1126/science.aax9552

54 Mehrabani, S., Bagherniya, M., Askari, G., Read, M. I. & Sahebkar, A. The effect of fasting or calorie restriction on mitophagy induction: a literature review. J Cachexia Sarcopenia Muscle 11, 1447–1458 (2020). 10.1002/jcsm.12611

55 Ölmez, T. T. et al. Sis2 regulates yeast replicative lifespan in a dose-dependent manner. Nature Communications 14, 7719–7719 (2023). 10.1038/s41467-023-43233-y

56 Li, L., Chen, O. S., McVey Ward, D. & Kaplan, J. CCC1 is a transporter that mediates vacuolar iron storage in yeast. J Biol Chem 276, 29515–29519 (2001). 10.1074/jbc.M103944200

57 Portnoy, M. E., Liu, X. F. & Culotta, V. C. Saccharomyces cerevisiae expresses three functionally distinct homologues of the nramp family of metal transporters. Mol Cell Biol 20, 7893–7902 (2000). 10.1128/MCB.20.21.7893-7902.2000

58 Biel, D., Steiger, T. K. & Bunzeck, N. Age-related iron accumulation and demyelination in the basal ganglia are closely related to verbal memory and executive functioning. Sci Rep 11, 9438 (2021). 10.1038/s41598-021-88840-1

59 Ndayisaba, A., Kaindlstorfer, C. & Wenning, G. K. Iron in Neurodegeneration - Cause or Consequence? Front Neurosci 13, 180 (2019). 10.3389/fnins.2019.00180

60 Cheng, R., Dhorajia, V. V., Kim, J. & Kim, Y. Mitochondrial iron metabolism and neurodegenerative diseases. Neurotoxicology 88, 88–101 (2022). 10.1016/j.neuro.2021.11.003

61 Fontana, L., Partridge, L. & Longo, V. D. Extending healthy life span--from yeast to humans. Science 328, 321–326 (2010). 10.1126/science.1172539

62 Kebbe, M., Sparks, J. R., Flanagan, E. W. & Redman, L. M. Beyond weight loss: current perspectives on the impact of calorie restriction on healthspan and lifespan. Expert Rev Endocrinol Metab 16, 95–108 (2021). 10.1080/17446651.2021.1922077

63 Sato, T., Shapiro, J. S., Chang, H. C., Miller, R. A. & Ardehali, H. Aging is associated with increased brain iron through cortex-derived hepcidin expression. Elife 11 (2022). 10.7554/eLife.73456

64 Kastman, E. K. et al. A Calorie-Restricted Diet Decreases Brain Iron Accumulation and Preserves Motor Performance in Old Rhesus Monkeys. Journal of Neuroscience 30, 7940–7947 (2010). 10.1523/JNEUROSCI.0835-10.2010

65 Xu, J., Knutson, M. D., Carter, C. S. & Leeuwenburgh, C. Iron Accumulation with Age, Oxidative Stress and Functional Decline. PLoS ONE 3, e2865–e2865 (2008). 10.1371/journal.pone.0002865

66 Cook, C. I. & Yu, B. P. Iron accumulation in aging: modulation by dietary restriction. Mech Ageing Dev 102, 1–13 (1998). 10.1016/s0047-6374(98)00005-0

67 Witte, A. V., Fobker, M., Gellner, R., Knecht, S. & Floel, A. Caloric restriction improves memory in elderly humans. Proc Natl Acad Sci U S A 106, 1255–1260 (2009). 10.1073/pnas.0808587106

68 Leclerc, E. et al. The effect of caloric restriction on working memory in healthy non-obese adults. CNS Spectr 25, 2–8 (2020). 10.1017/S1092852918001566

69 Goudswaard, L. J. et al. Using trials of caloric restriction and bariatric surgery to explore the effects of body mass index on the circulating proteome. Scientific Reports 13, 21077–21077 (2023). 10.1038/s41598-023-47030-x

70 Capel, F. et al. Macrophages and Adipocytes in Human Obesity. Diabetes 58, 1558–1567 (2009). 10.2337/db09-0033

71 Siino, V. et al. Plasma proteome profiling of healthy individuals across the life span in a Sicilian cohort with long-lived individuals. Aging Cell 21 (2022). 10.1111/acel.13684

72 Lehallier, B. et al. Undulating changes in human plasma proteome profiles across the lifespan. Nature Medicine 25, 1843–1850 (2019). 10.1038/s41591-019-0673-2

73 Sebastiani, P. et al. Protein signatures of centenarians and their offspring suggest centenarians age slower than other humans. Aging Cell 20 (2021). 10.1111/acel.13290

74 Pagani, A., Nai, A., Silvestri, L. & Camaschella, C. Hepcidin and Anemia: A Tight Relationship. Front Physiol 10, 1294 (2019). 10.3389/fphys.2019.01294

75 Milic, S. et al. The Role of Iron and Iron Overload in Chronic Liver Disease. Med Sci Monit 22, 2144–2151 (2016). 10.12659/msm.896494

76 Gammella, E. et al. Unconventional endocytosis and trafficking of transferrin receptor induced by iron. Mol Biol Cell 32, 98–108 (2021). 10.1091/mbc.E20-02-0129

